# Biofabrication of spatially organized temporo-mandibular fibrocartilage assembloids

**DOI:** 10.1101/2024.12.20.629744

**Authors:** Alexandre Dufour, Lucie Essayan, Beomjoon Kim, Vincent Salles, Christophe Marquette

## Abstract

The combination of mesenchymal stem cell (MSC) spheroids and polymeric scaffolds has been actively explored for engineering organized hyaline cartilage; however, its application to other types of cartilage remains under-explored. The temporo-mandibular joint (TMJ) fibrocartilage is a highly stratified tissue whose recapitulation remains challenging. In this study, the shape and growth orientation of assembloids were controlled by seeding early mature human adipose-derived MSC spheroids into polymeric scaffolds with a dual architecture of micron-scale fibers. This results in flattened asymmetric tissues with a single-sided articular surface. Structurally, the engineered fibrocartilage mimicked the histotypical organization observed in the native human condylar fibrocartilage, notably featuring a thick fibrous zone with flattened cells. Native-like distribution of general extracellular matrix (ECM) components, including glycosaminoglycans and total collagens, and ECM-specific components, such as type I and II collagens, aggrecan core protein, and fibronectin, were observed. Collagen organization, as demonstrated by polarized light microscopy and scanning electron microscopy at the fibril level, was also found to be similar to that of native human tissue. Zonal-dependent micromechanical properties were identified in both the engineered and native tissues, although lower mechanical properties were observed in the fibrous zone of the engineered tissue. This work provides further evidence that the combination of MSC spheroids and micron-sized fiber polymeric scaffolds is a versatile approach for engineering stratified cartilage and a promising strategy for engineering biomimetic fibrocartilage grafts for TMJ reconstruction.

## Introduction

The temporo-mandibular joint (TMJ) is an articulation that links the upper and lower jaws by interfacing the mandible and the temporal bone. It is a paired or bicondylar joint, with each joint formed by the glenoid fossa and articular eminence of the temporal bone, the condylar head of the mandible, and the interposed articular disc separating the joint in two compartments [1]. It is a ginglymoarthrodial joint, with "ginglymus" referring to its hinge joint properties, allowing motion only backward and forward in one plane, and "arthrodial" referring to the gliding of the surfaces. The combination of hinging and sliding enables various movements to occur simultaneously at both condylar heads, which in turn facilitates essential functions, such as chewing, speaking, and breathing [2].

The articular surfaces of both condylar heads and glenoid fossa are distinguishable from most other articulations because they are covered by fibrocartilage instead of the typical hyaline cartilage. The outline of TMJ articular surfaces can change in certain temporo-mandibular disorders (TMDs), such as TMJ osteoarthritis (OA), which is primarily degenerative and leads to deterioration of the articular surface, resulting in pain and dysfunction of the joint. The most commonly affected surface is the mandibular condyle, more rarely, these changes involve the articular surface of the glenoid fossa [3]. TMJ-OA affects between 8 % and 16 % of the population, whereas knee-OA affects over 14 % [4], with the primary cause assumed to be excessive mechanical loading on the articular fibrocartilage [5]. As TMJ-OA progresses, common clinical symptoms include orofacial pain, limited mouth opening, and joint clicking sounds, which affect the aforementioned essential functions [6].

When conservative treatments fail or when most of the TMJ anatomical structures are impaired, surgical intervention is the leading treatment option. Functional restoration can be performed using autologous chondro-costal grafts, free fibula, or vascularized medial femoral condyle flaps; however, additional lesions and risks of complications at the donor site are major drawbacks [7,8]. Alternatively, TMJ replacement with a synthetic prosthesis that combines polymers and alloys has shown reasonable outcomes, with some studies reporting over 90 % success rates, including decreased pain and increased maximal opening up to 20 years post-surgery [9,10]. Approximately 450 patients per year underwent this procedure from 2005 to 2014 in the US, and a 30 % increase is projected by 2030 [11]. Nevertheless, postoperative complications such as metal particulation leading to osteolysis have been reported, posing a risk for revision surgery [12]. Additionally, metal hypersensitivity and nickel allergies have raised concerns among surgeons, with up to 17 % of individuals having metal allergies. Metal hypersensitivity to an implanted joint prosthesis can manifest as swelling, pain, joint effusions, and prosthesis failure [13,14]. A tissue engineering approach may address the issues of donor site morbidity, allergy, and limited longevity by generating autologous viable fibrocartilage tissue capable of self-renewal and normal function, providing surgeons with the biological material to completely regenerate damaged TMJ structures.

Cartilage has a complex composition and structural organization, which is difficult to replicate in engineered tissues. Recent studies focusing on the engineering of hyaline cartilage for hip or knee replacement have made significant progress in bridging this gap by creating biomimetic tissues that mimic the organization of the collagen network. This was achieved through the guided growth of self-assembled mesenchymal stem cell (MSC) spheroids within 3D-printed polymeric scaffolds, mimicking the process of mesenchymal condensation, so that cartilage tissues form in a manner reminiscent of cartilage morphogenesis [15,16]. The physical constraints provided by these scaffolds guided the formation of a collagen network by the cells. This approach was initially demonstrated using large polymeric fibers (> 200 μm) produced through fused deposition modelling [17] and has since been extended to micron-scale fibers created by melt-electrowriting [18]. This biofabrication strategy has been shown to produce a collagen network that mimics developmental-like organization, with a Benninghoff-like orientation of collagen fibrils and mechanical behavior similar to that of the native tissue.

The TMJ condyle also develops from MSC condensation, namely the condylar blastema, which differentiates into condylar cartilage. This cartilage then undergoes endochondral ossification, similar to the process in the long bones [19]. Given this developmental process, the use of MSC spheroids is also of interest for engineering TMJ fibrocartilage. The latter exhibits specific depth-dependent variations in extracellular matrix (ECM) composition and organization, which are crucial for the normal function of the condyle. From the articular surface to the subchondral bone, TMJ fibrocartilage is composed of four distinct zones: fibrous, proliferative, mature, and hypertrophic zones, with type I collagen present throughout all layers. The fibrous zone contains flattened fibrochondrocytes embedded within a dense ECM of type I collagen fibers oriented parallel to the articular surface. The proliferative zone is primarily a cellular region characterized by a relatively high proportion of type I collagen. The mature and hypertrophic zones resemble hyaline cartilage found in other joints, with chondrocytes as the predominant cell type within an ECM that primarily consists of randomly organized type II collagen [20].

In this study, we built on recent advances in the engineering of biomimetic hyaline cartilage to develop fibrocartilage. Our approach involved assembling human MSC multicellular spheroids into polymeric scaffolds with a dual architecture of micron-scale fibers to engineer fibrocartilage assembloids. We hypothesized that the constraints provided by the physical boundaries of the scaffold, combined with the initial state of the cells (i.e., pre-formed instead of self-assembled within the scaffold), would facilitate native-like cell morphology, collagen organization, and ECM deposition. We aimed to show that the combination of scaffold constraints with pre-assembled MSC spheroids would result in native-like stratified fibrocartilage tissue to address the unmet clinical need for functional fibrocartilage grafts.

## Results

### 1. Direct-Writing Electrospinning (DWE)

To capture, culture, and orient the growth of cell spheroids, scaffolds consisting of two types of meshes were produced by extruding solubilized poly(ε-caprolactone) (PCL) through an electric field. The primary mesh was a box-like structure measuring 19 × 19 mm, with each fibrous layer oriented at 90° to the previous layer and spaced 0.9 mm apart. Additionally, a secondary mesh measuring 23 mm × 23 mm, referred to as catching fibers [21], was included. Each fibrous layer of this mesh was also oriented at 90° to the previous layer, with a spacing of 0.15 mm, resulting in a usable area with a 10 mm diameter (**Figure 1-Ai**). The printing order consisted of five layers of the primary mesh, followed by three layers of catching fibers at a 45° angle and 295 layers of the primary mesh (**Figure 1-Aii**). After fine-tuning the nozzle-to-collector distance in increments of 30 µm for every 50 printed layers (**Supplemental Figure 1**) and the timing of the catching fiber layer deposition, as described above (**Supplemental Figure 2**), scaffolds matching the design specifications were successfully produced in approximately 45 min (**Figure 1-Bi, Bii, Biii, and Biv**). Qualification of the DWE process through scanning electron microscopy (SEM) (**Figure 1-Ci, Cii, Ciii, and Civ**) and digital microscopy (**Supplemental Figure 3**) demonstrated a linear correlation between the number of deposited layers and scaffold thickness, resulting in a total scaffold thickness of 1 mm for the specified design. SEM measurements also showed that the fibers forming the primary mesh had an average diameter of 4.7 ± 0.4 µm, with a spacing of 942 ± 34 µm at the bottom and 956 ± 51 µm at the top of the structure. In contrast, the fibers in the catching fiber network averaged 13.8 ± 3.1 µm in diameter with a spacing of 144 ± 71 µm.

**Figure 1.**
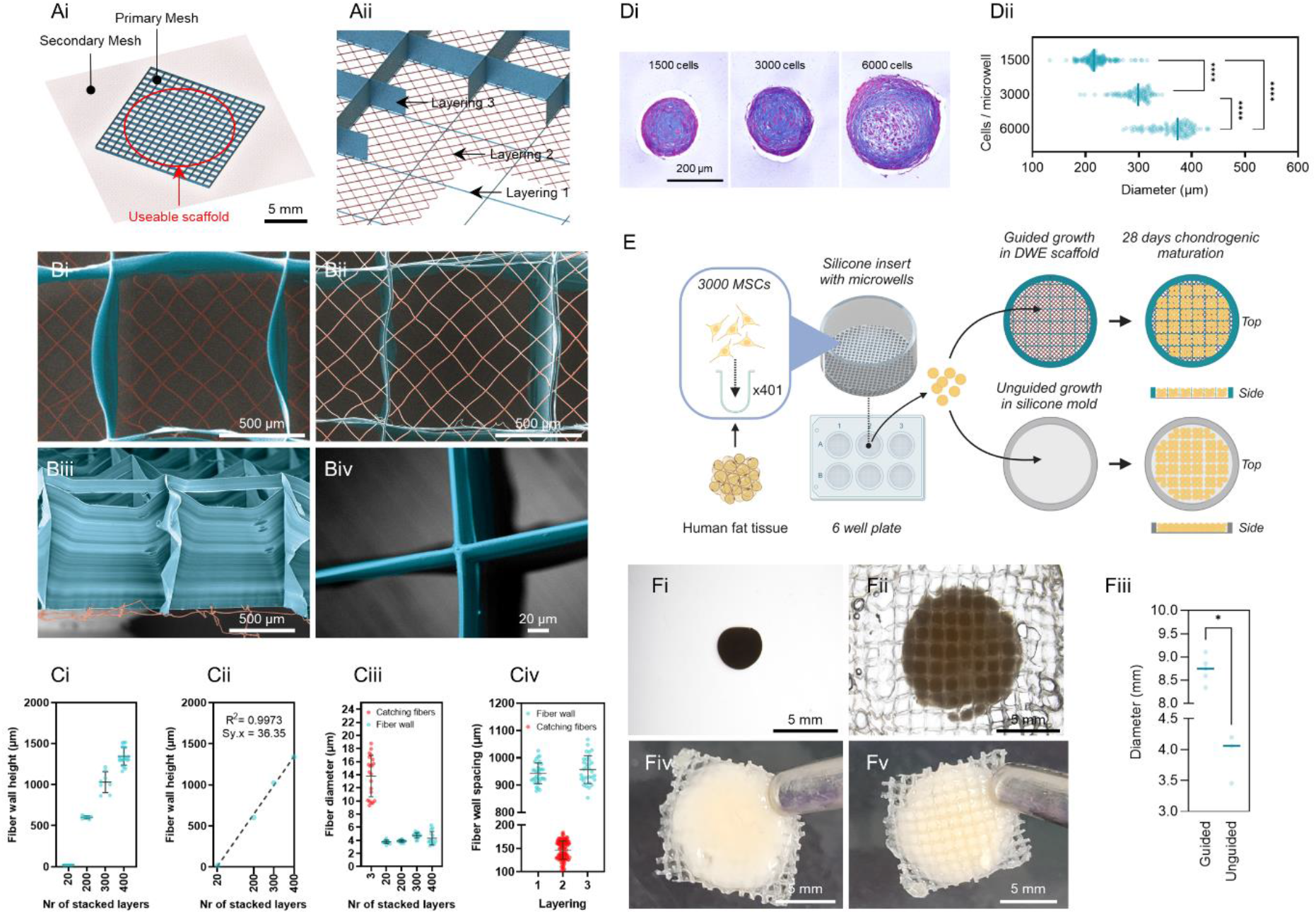
Direct-writing electrospinning (DWE) of multiphasic scaffold and construction of fibrocartilage assembloids. (Ai, Aii) Schematics of (Ai) design and (Aii) printing strategy of a dual- sized scaffold with a primary mesh (blue, 900 µm spacing) and a secondary mesh (red, 150 µm spacing) of 45° angled and closely spaced fibers to retain spheroids. (Bi-Biv) Scanning electron microscopy (SEM) images of printed scaffold with the main mesh and the secondary mesh false- colored in blue and red respectively. The scaffold is viewed from (Bi & Biv) the top, (Bii) the bottom, and (Biii) the cross-section. (Ci, Cii) Fiber wall height, (Cii) fiber diameter and (Civ) fiber wall spacing measured from SEM images. The coefficient of determination (R^2^) and the standard deviation of the residuals (Sy.x) are indicated in (Cii). (Di) Human adipose-derived mesenchymal stem cells spheroids with different initial cell densities cultured for a week in chondrogenic medium and stained for alcian blue, with (Dii) corresponding spheroid diameters. (E) A schematic of the workflow experiment used for generating fibrocartilage assembloids. (Fi-Fv) Stereomicroscopic images of (Fi) guided and (Fii) unguided fibrocartilage assembloids after 4 weeks of chondrogenic culture, with (Fiii) corresponding diameters. (Fiv, Fv) Macroscopic images of guided fibrocartilage assembloids from (Fiv) top (articular surface) and (Fv) bottom (spheroid-catching fiber side) after 4 weeks of chondrogenic culture.

### 2. Biofabrication and Macroscopic Aspect of Fibrocartilage Assembloids

Fibrocartilage assembloids were generated using human adipose-derived (AD) MSC spheroids, each composed of 3,000 cells, representing the largest histologically uniform cell aggregate (294 ± 28 µm) that could be formed under the conditions investigated (**Figure 1-Di and Dii**). These spheroids were produced at a high throughput in chondrogenic media using custom silicone microwells. To study the influence of structural guidance on tissue growth, two conditions were established: guided engineered tissue, in which spheroids were seeded into DWE scaffolds, and unguided engineered tissue, in which spheroids were placed in a control silicone mold designed to produce unoriented tissue growth. AD-MSCs spheroids were seeded into the DWE scaffolds or control silicone mold 48 h after their formation (**Figure 1-E**). The seeded volume for both conditions was a cylinder with a diameter of 10 mm and height of 1 mm (i.e., the height of the DWE scaffold). Based on this volume and the size of the spheroids [22], a total of 4,468 spheroids were seeded per structure (≈ 57 spheroids/mm³).

In the absence of a polymeric mesh, the AD-MSC spheroids contracted dramatically, forming spherical tissues (**Figure 1-Fi**). DWE scaffolds prevented this contraction, resulting in flattened constructs (**Figure 1-Fii**) that were approximately twice the diameter of the unguided tissues (**Figure 1-Fiii**). Notably, significant tissue growth was observed toward the top of the scaffold (**Figure 1-Fiv and Fv**), where catching fibers were absent. This tissue development, referred to as the articular surface, demonstrates architecture-guided tissue growth.

### 3. Histomorphometry and Overall Distribution and Quantification of ECM

Several classification schemes describe the zonal organization of the fibrocartilage in the mandibular condyle. The most common is the four-zone nomenclature, which includes the fibrous, proliferative, mature, and hypertrophic zones. The fibrous zone corresponds to the articular surface and is separated from the underlying hyaline-like mature and hypertrophic zones by a highly cellular proliferative zone [20]. In this study, we used a three-zone nomenclature that refers to the fibrous (FZ), proliferative (PZ), and hyaline-like (HZ) zones. The latter combines the mature and hypertrophic zones, which share characteristics resembling the hyaline cartilage found in other joints.

Measurements of aspect ratio and Feret’s diameter on haematoxylin and eosin (H&E) stained tissue sections confirmed that unguided tissues were spherical (1.6 ± 0.006) with a smaller diameter (3.2 ± 0.37 mm), while guided tissues exhibited a significantly flattened profile (7.8 ± 0.007) and a larger diameter (7.7 ± 0.76 mm) (**Figure 2-A** and **Figure 2-G and H**). Histological comparison of engineered tissues with human adult mandibular condyle fibrocartilage revealed that both displayed a FZ on the surface, characterized by flattened cells and a layered tissue structure.

**Figure 2.**
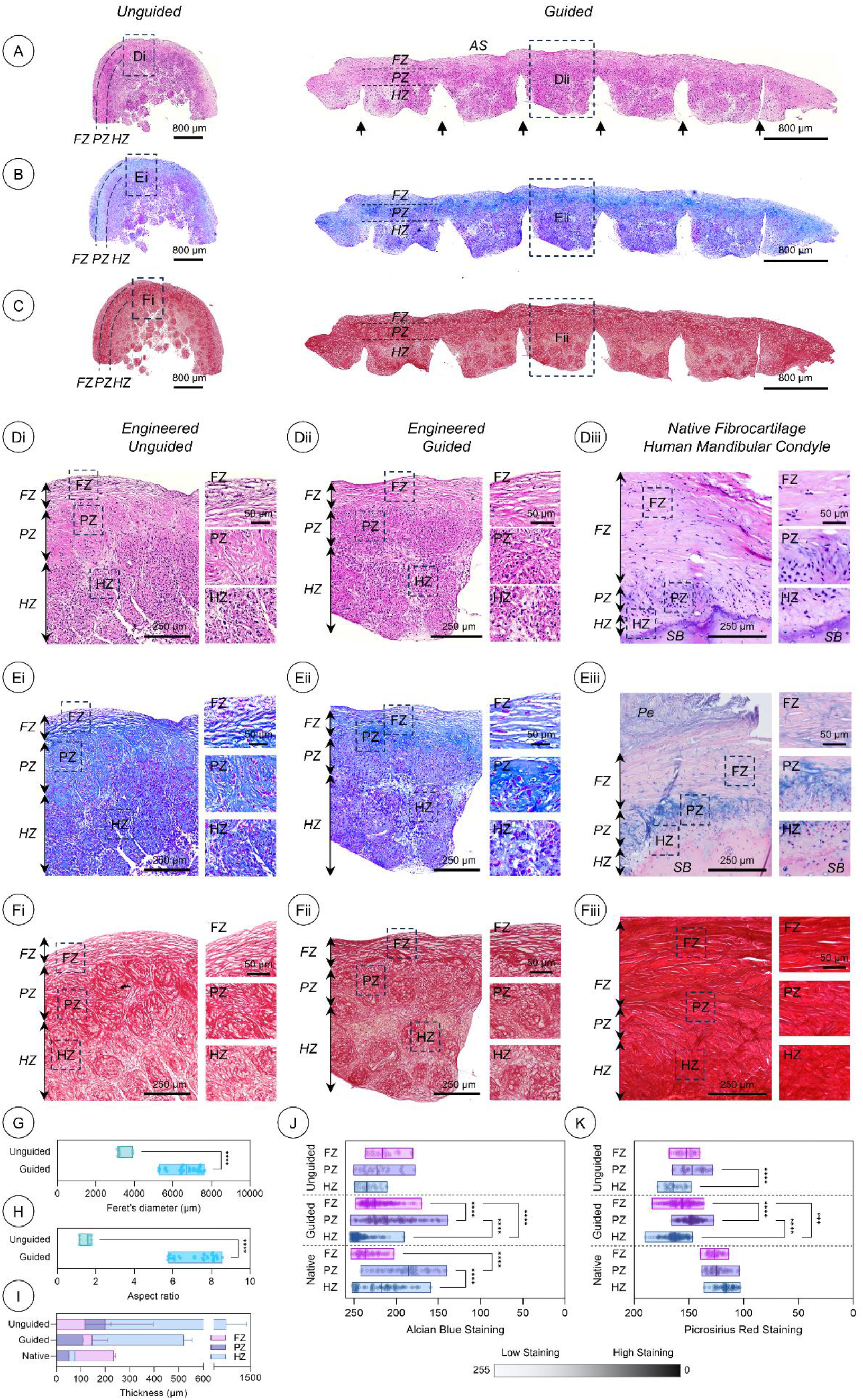
General morphology and extracellular matrix distribution in engineered and native fibrocartilage. (A-C) Plain histological sections of engineered tissues stained with (A) haematoxylin and eosin (H&E) for general morphology, (B) alcian blue and nuclear fast red for glycosaminoglycans (GAG), and (C) picrosirius red for collagen content. Black arrows indicate the position of the polymeric fiber walls (dissolved during histological processing). (Di-Fiii) High magnification views of (D) H&E, (E) alcian blue and nuclear fast red, and (F) picrosirius red staining in (Di, Ei, Fi) unguided, (Dii, Eii, Fii) guided engineered tissues, and (Diii, Eiii, Fiii) native human mandibular condyle fibrocartilage. (G-I) Histomorphometric measurements in H&E-stained sections. (J, K) Quantitative analysis of (J) alcian blue after color deconvolution, and (K) picrosirius red staining. Data are represented as (G, H, J, K) bars with minimum, maximum, and median values or (I) as mean with error bars. Each single data point represents (G, H) a tissue section or (J, K) a distinct histological region. Multiple tissue sections and histological regions were analyzed per sample (unguided, n = 3; guided, n = 4; native, n = 1). ***, **** denote statistical significance with P < 0.001 and P < 0.0001, respectively (Mann-Whitney test for two groups, or Kruskal-Wallis test followed by Dunn’s multiple comparisons test). AS, articular surface ; FZ, fibrous zone ; PZ, proliferation zone ; HZ, hyaline zone ; Pe, perichondrium.

In the unguided tissue, the FZ extended across the entire surface of the spherical tissue. H&E staining revealed two distinct regions beneath the FZ, referred to as the PZ and HZ, moving inward from the surface (**Figure 2-Di**). In the guided tissues, notable tissue growth was observed on the top side, featuring a FZ followed by an additional region located between the FZ and the polymeric mesh. This adjacent region was categorized as the PZ. The remaining tissue bordered by the catching fibers was referred to as the HZ (**Figure 2-Dii**). In the native TMJ, the FZ was bordered by the perichondrium at the top and the more densely cellularized PZ at the bottom. Following this, the HZ was ultimately bound by the subchondral bone (**Figure 2-Diii**). Zonal measurements indicated that the FZ comprised the majority of the native tissue section (249 ± 270 µm), followed by the HZ (78 ± 34 µm) and PZ (59 ± 31 µm) (**Figure 2-I**). In engineered tissues, the HZ occupied most of the section (1171 ± 136 µm for unguided and 514 ± 61 µm for guided tissues, respectively). In unguided tissue, this was followed by PZ (191 ± 72 µm) and FZ (104 ± 46 µm). In the guided tissues, HZ was followed by FZ (145 ± 51 µm) and PZ (118 ± 60 µm).

Both unguided and guided tissues exhibited positive glycosaminoglycan (GAG) deposition, as evidenced by alcian blue staining, with distinct staining patterns consistently observed throughout the tissue sections (**Figure 2-B**). In unguided tissues, the staining intensity progressively decreased from the surface to the deeper layers. In contrast, guided tissues displayed maximum staining beneath the FZ, specifically at the PZ level, similar to the native tissue (**Figure 2-Ei, Eii, and Eiii**). This pattern of GAG deposition was consistently observed in disease-free portions of the native human condylar fibrocartilage (**Supplemental Figure 4**), which served as a control throughout the study and was free of defects and vessel invasion (**Supplemental Figure 5**). This GAG deposition pattern was more clearly observed after color deconvolution of alcian blue and nuclear fast red-stained tissue sections (**Supplemental Figure 6**) and was used to quantify the alcian blue staining intensity. This confirmed a significant differential in zonal GAG deposition in both guided and native tissues, supporting the similar deposition patterns seen in both (**Figure 2-J**). The presence of GAGs in engineered tissues was further demonstrated through biochemical quantification (**Table 1**) (**Supplemental Figure 7**), which revealed no significant difference between the engineered and native fibrocartilage.

**Table 1.** Biochemical properties of engineered and native fibrocartilage. GAG, DNA, and total collagen (OHP) contents were normalized to the wet weight (wt.). *, ** and *** denote a significant difference when compared to native tissue with P < 0.05, P < 0.01 and P < 0.001, respectively (ANOVA followed by Kruskal-Wallis multiple comparisons).

| | | DNA/Wet wt. ($\mu\text{g}/\text{mg}$) | GAG/Wet wt. ($\mu\text{g}/\text{mg}$) | OHP/Wet wt. ($\mu\text{g}/\text{mg}$) |
| --- | --- | --- | --- | --- |
| Engineered<br>Unguided | Mean ± Std Dev. | $1.84 \pm 0.4$ | $4.99 \pm 2.33$ | $35 \pm 5.93$ |
|  | Median ; Range | 1.98 ; 0.8 | 4 ; 4.33 | 34.5 ; 13 * |
| Engineered<br>Guided | Mean ± Std Dev. | $2.26 \pm 1$ | $4.15 \pm 1.26$ | $30.92 \pm 12.02$ |
|  | Median ; Range | 2.3 ; 2.1 ** | 4.5 ; 4.08 | 32.46 ; 49.23 *** |
| Native | Mean ± Std Dev. | $0.81 \pm 0.33$ | $3.55 \pm 0.74$ | $291 \pm 52.95$ |
|  | Median ; Range | 0.74 ; 0.93 | 3.42 ; 1.65 | 291.48 ; 161.17 |

Collagen deposition was also evidenced by positive picrosirius red staining (**Figure 2-C**). The staining intensity for total collagen deposition was notably stronger in the PZ of engineered tissues, whereas it remained relatively homogeneous throughout the native fibrocartilage (**Figure 2-Fi, Fii, and Fiii**). This observation was quantitatively confirmed, showing an overall stronger staining intensity in native tissues than in engineered tissues (**Figure 2-K**). Biochemical quantification further revealed that the collagen content in the native fibrocartilage was ten times higher than that in the engineered tissues (**Table 1**) (**Supplemental Figure 7**).

### 4. Nuclei Distribution, Morphology, and Cell Proliferative Capacity

Nuclei distribution and morphology varied across tissue sections in both native and engineered tissues (**Figure 3-A**). A lower density of flattened nuclei was observed in the FZ of guided tissues (1,327 ± 469 nuclei/mm² with an aspect ratio of 2 ± 0.9), while the PZ (4,985 ± 765 nuclei/mm²; 1.6 ± 0.6) and HZ (5,176 ± 798 nuclei/mm²; 1.6 ± 0.5) exhibited similar densities of oval nuclei. Similar results were observed in the unguided tissues (**Supplemental Table 1**). In the native tissue, a low density of flattened nuclei was also found in the FZ (735 ± 229 nuclei/mm²; 2 ± 0.86), with numerous oval nuclei in the PZ (1,623 ± 479 nuclei/mm²; 1.78 ± 0.64). The HZ displayed similar nuclei density to the FZ, but with oval nuclei (619 ± 201 nuclei/mm²; 1.68 ± 0.65). Quantitative analysis indicated similar cell densities in the FZ of both the engineered and native tissues. However, the engineered tissues exhibited a greater number of nuclei in the PZ and HZ (**Figure 3-B**). This finding was corroborated by DNA quantification, which revealed that the engineered tissues contained twice the DNA content of native tissues (**Table 1**) (**Supplemental Figure 7**). In terms of nuclei morphology, the distribution patterns were comparable between tissue types (**Figure 3-C** and **Figure 3-D**).

**Figure 3.**
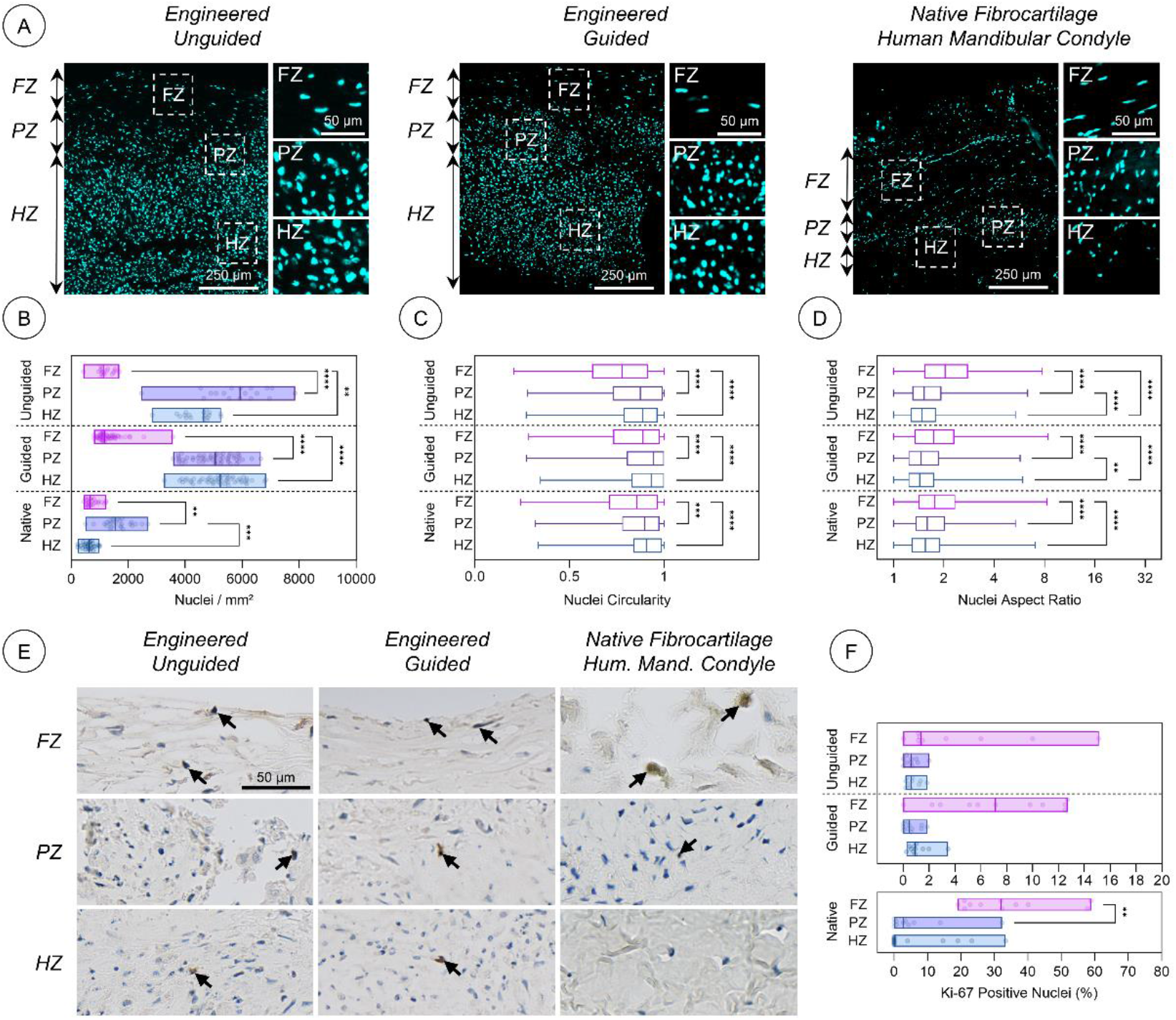
Nuclei distribution, morphology and cell proliferative capacity. (A) Histological sections of engineered and native tissues with nuclei stained in cyan. (B-D) Zone-dependent (B) density and (C, D) morphology of nuclei. (E) Ki-67 immunostaining to assess cell proliferation in engineered and native tissues. Black arrows indicate Ki-67 positive nuclei. (F) Quantification of Ki-67 positive nuclei in the histological regions. Data in panels (B) and (F) are shown as bars representing minimum, maximum, and median values, with each data point corresponding to a unique histological region. Panels (C) and (D) display data in a box-and-whisker format. Multiple sections and histological regions were analyzed per sample (unguided, n = 3; guided, n = 4; native, n = 1). **, ***, and **** denote statistical significance at P < 0.01, P < 0.001, and P < 0.0001, respectively (Kruskal-Wallis test followed by Dunn’s multiple comparisons test). FZ, fibrous zone ; PZ, proliferation zone ; HZ, hyaline zone.

Evaluation of the cell proliferative capacity in tissue sections revealed that proliferative cells were primarily located in the FZ of the tissues (**Figure 3-E**). In the native fibrocartilage, proliferative cells were not detected in the HZ and were ten times lower in the PZ (≈ 3 %) than in the FZ (≈ 32 %). (**Figure 3-F**). A similar trend was observed in engineered tissues, although to a lesser extent (≈ 1.4 %, 0.6 %, and 0.6 % for unguided tissue and 7 %, 0.5 %, and 0.9 % for guided tissue in the FZ, PZ, and HZ, respectively).

### 5. Spatial Distribution of ECM-specific Tissue Components

The presence and spatial distribution of type I and II collagens, along with the aggrecan core protein and fibronectin, were assessed by immunofluorescence in both engineered and native tissues (**Figure 4-A**) (**Supplemental Figure 8** and **Supplemental Figure 9**). Type I collagen, the main component of fibrocartilage, was strongly detected throughout the tissue sections of both the engineered and native fibrocartilage. Overall, quantification of type I collagen immunofluorescence showed a significantly higher level of staining in native fibrocartilage than in engineered tissues (**Figure 4-Bi**). In the native tissue, a decrease in type I collagen immunofluorescence was observed from the FZ to the HZ. The opposite trend was noted in the guided tissue, with twice as much signal detected in the HZ compared to the FZ (**Figure 4-Bii**), while no zonal difference was detected in the unguided tissue. However, none of these trends was statistically significant.

**Figure 4.**
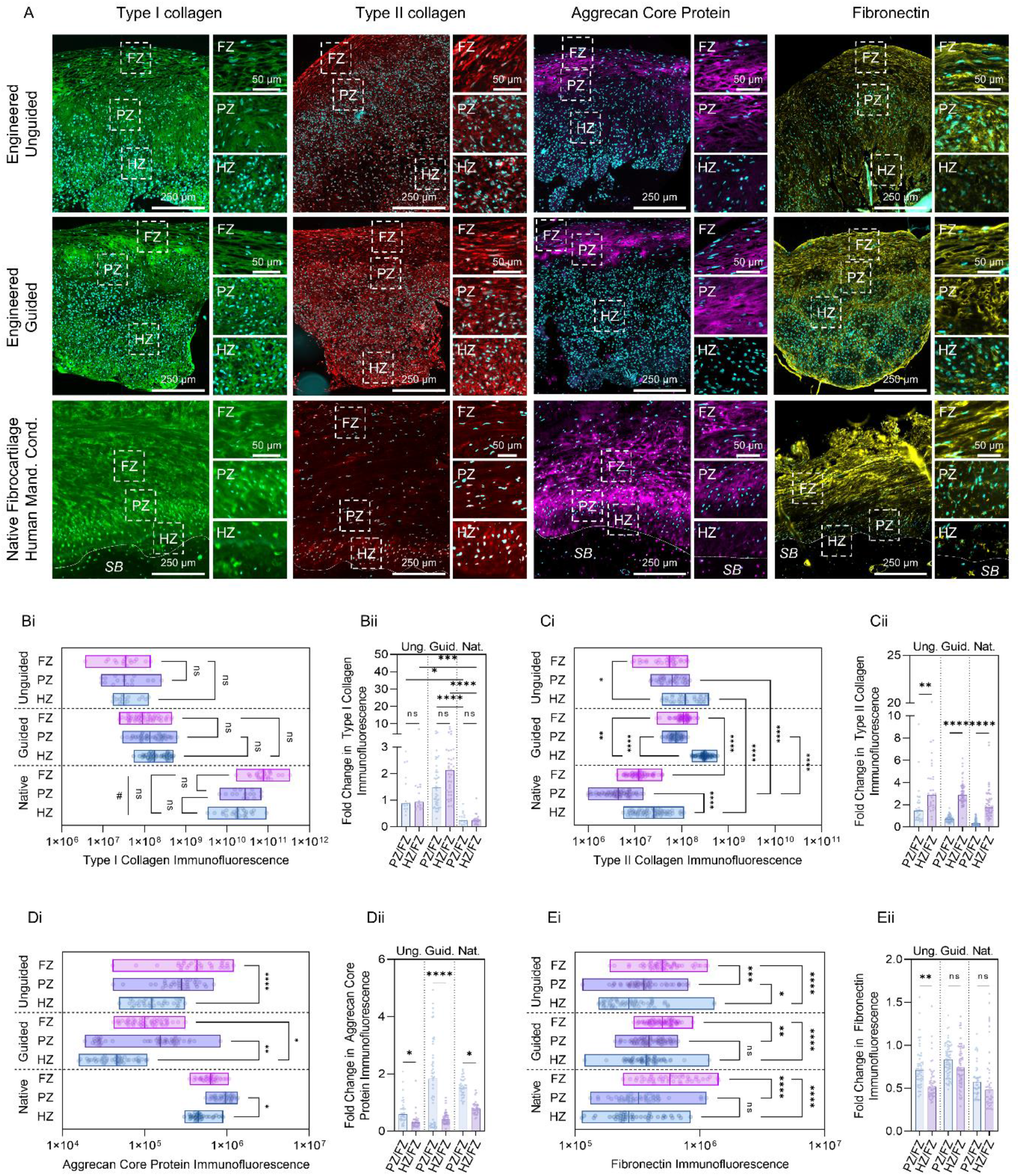
Distribution of extracellular matrix-specific components in engineered and native fibrocartilage. (A) Immunofluorescence staining for type I (green) and II (red) collagens, aggrecan core protein (purple), and fibronectin (yellow). The dotted lines represent the frontier between the hyaline zone and the subchondral bone in native tissue. (Bi, Ci, Di, Ei) Quantitative evaluation of zonal immunofluorescence and (Bii, Cii, Dii, Eii) zonal fold change of (Bi, Bii) type I collagen, (Ci, Cii) type II collagen, (Di, Dii) aggrecan core protein, and (Ei, Eii) fibronectin. Data in panels (Bi, Ci, Di, Ei) are shown as bars representing minimum, maximum, and median values. Data in panels (Bii, Cii, Dii, Eii) are displayed as bars indicating the median. Each single data point represents a distinct histological region. Multiple histological regions were analyzed on multiple tissue sections per sample (unguided, n = 3; guided, n = 4; native, n = 1). *, **, ***, **** denote statistical significance with P < 0.05, P < 0.01, P < 0.001, and P < 0.0001, respectively (Kruskal-Wallis test followed by Dunn’s multiple comparisons test). FZ, fibrous zone ; PZ, proliferation zone ; HZ, hyaline zone ; SB, subchondral bone.

Type II collagen was also found throughout the tissue sections in both engineered and native fibrocartilage, but with lower level of staining in native fibrocartilage than in engineered tissues (**Figure 4-Ci**). Notably, there were significant zonal variations in type II collagen deposition. In human fibrocartilage, HZ showed the strongest type II collagen signal, with nearly double the intensity compared to FZ. This was followed by FZ and PZ, which exhibited similar staining intensity. Interestingly, guided engineered tissue exhibited the same zonal variation in type II collagen distribution as native fibrocartilage, while unguided tissue displayed a more linear transition in staining intensity, with HZ showing the strongest signal (**Figure 4-Cii**).

Aggrecan core protein also demonstrated significant zonal variations, with a trend indicating higher staining levels in native fibrocartilage than in engineered tissues (**Figure 4-Di**). The PZ exhibited the highest signal intensity in the native tissue, nearly double that of the FZ, while the HZ and FZ displayed similar staining levels (**Figure 4-Dii**). Guided engineered tissue reflected the same zonal variation as native fibrocartilage, whereas unguided tissue showed a more linear transition in staining intensity, with the strongest signal in the FZ.

Finally, fibronectin exhibited a linear transition in staining intensity in both engineered and native tissues, with the strongest signal observed in the FZ. This zonal variation was significant across all zones in unguided tissues (**Figure 4-Ei**). In contrast, in both native and guided tissues, only the FZ exhibited significant variations, with signal intensity at least double that of PZ and HZ. (**Figure 4-Eii**).

### 6. Organization of the Collagen Network Through Polarized Light Microscopy (PLM)

The overall architecture of the collagen network was first investigated using PLM of tissue sections from both engineered and native tissues (**Figure 5-A**) (**Supplemental Figure 10**). Collagen fibrils were predominantly oriented parallel to the surface in the FZ, whereas the PZ and HZ exhibited randomly oriented bundles in both engineered and native tissues (**Figure 5-Bi, Bii, Biii**). Color-coding of PLM images confirmed these observations (**Figure 5-C**), highlighting the differential zonal orientation in both engineered and native tissues (**Figure 5-Di, Dii, Diii**). Quantitatively, the pattern of fibril direction was highly similar between the guided and native tissues (6 ± 6 °, 17 ± 16 °, 40 ± 28 ° and 6 ± 5 °, 19 ± 10 °, 38 ± 13 ° for FZ, PZ and HZ respectively) (**Figure 5-E**), but notable zonal differences in fibril dispersion (**Figure 5-F**) and alignment quality (**Figure 5-G**) were primarily observed in the guided tissues (**Supplemental Table 2**). Nevertheless, the overall trends were consistent between engineered and native tissues.

**Figure 5.**
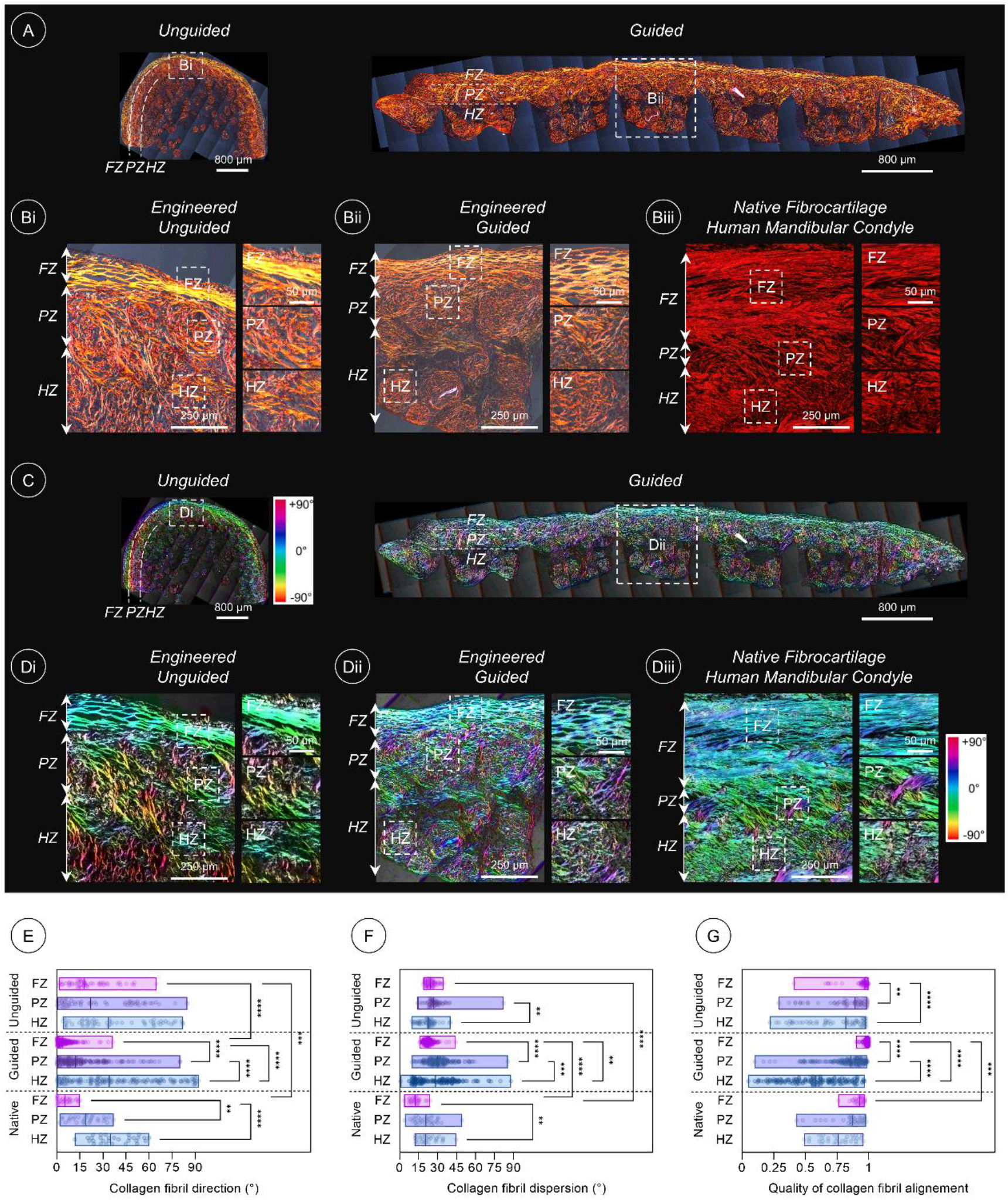
Orientation of collagen network analyzed through polarized light microscopy (PLM). (A, Bi-Biii) Original PLM images. (A) Plain histological sections of engineered tissues. (Bi-Biii) High- magnification views of (Bi) unguided and (Bii) guided engineered tissues, and (Biii) native human mandibular condyle fibrocartilage. (B, Ci-Ciii) Color-coded PLM images. (E-G) Quantitative evaluation of (E) collagen fibril orientation, (F) dispersion, and (G) quality of alignment. Data are represented as bars showing minimum, maximum, and median values. Each single data point represents a distinct histological region. Multiple histological regions were analyzed on multiple tissue sections per sample (unguided, n = 3; guided, n = 4; native, n = 1). **, ***, and **** denote statistical significance at P < 0.01, P < 0.001, and P < 0.0001, respectively (Kruskal-Wallis test followed by Dunn’s multiple comparisons test). FZ, fibrous zone ; PZ, proliferative zone ; HZ, hyaline zone.

### 7. Scanning Electron Microscopy (SEM) Analysis of Tissues and Collagenic ECM

SEM analysis of human mandibular condyle fibrocartilage (**Figure 6-A**) revealed a smooth articular surface and zonal organization characterized by tissue layering in the FZ with aligned tissue fibers (**Figure 6-B**). In contrast, the tissues in the PZ and HZ were randomly organized. The guided engineered tissue displayed a similar appearance and organization, with tissue growth confined by catching fibers (**Figure 6-C**). Additionally, a transition in cell morphology was observed from flat cells in the FZ to spherical cells in the PZ and HZ (**Supplemental Figure 11**). This overall structure was maintained even after the removal of non-collagenous components, suggesting the stratification of the collagen network in the engineered tissue along distinct planes (**Figure 6-D**). Investigations at high magnification in guided engineered tissue revealed a similar alignment of tissue fibers in the FZ to that in native tissue [23], with parallel stacking of tissue layers (**Figure 7**). In contrast, the fibers were randomly arranged in the PZ and HZ. Similar observations were made for collagen fibril bundles with an unobstructed view of the collagen network, demonstrating native-like collagen fibril organization.

**Figure 6.**
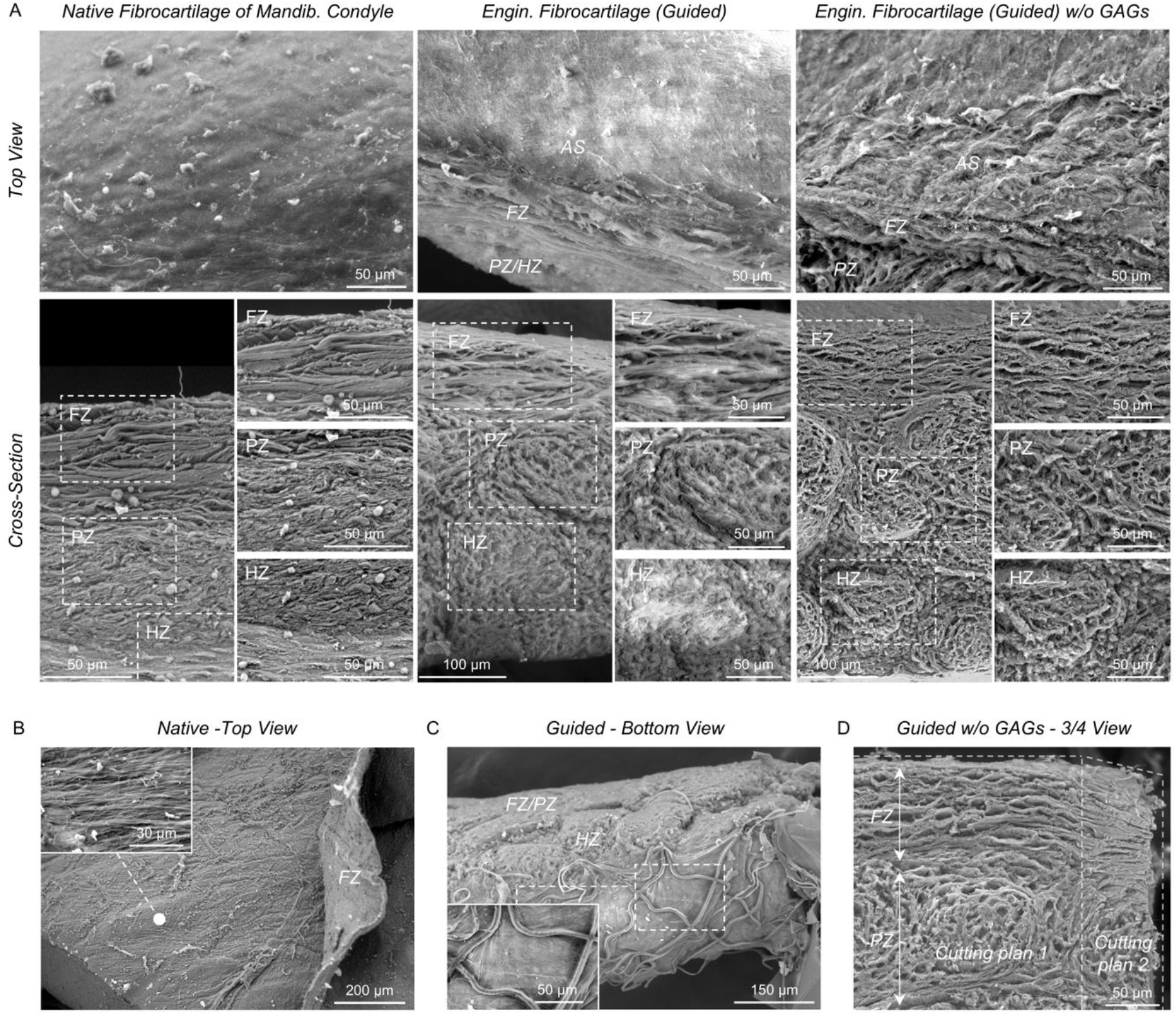
Scanning electron microscopy observations of guided engineered and native fibrocartilage. (A) Top and cross-sectional views of the human mandibular condyle fibrocartilage (left panel), guided engineered fibrocartilage (middle panel), and guided engineered fibrocartilage after glycosaminoglycan removal (w/o GAGs) by serial enzymatic digestion (right panel). (B) Top view of human mandibular condyle fibrocartilage with the fibrous zone peeled off, revealing native fibril orientation. (C) Bottom view of guided engineered fibrocartilage showing the bottom tissue and catching fibers. (D) ¾ view of GAG-depleted guided engineered fibrocartilage showing collagen organization across two planes. AS, articular surface; FZ, fibrous zone; PZ, proliferative zone; HZ, hyaline zone.

**Figure 7.**
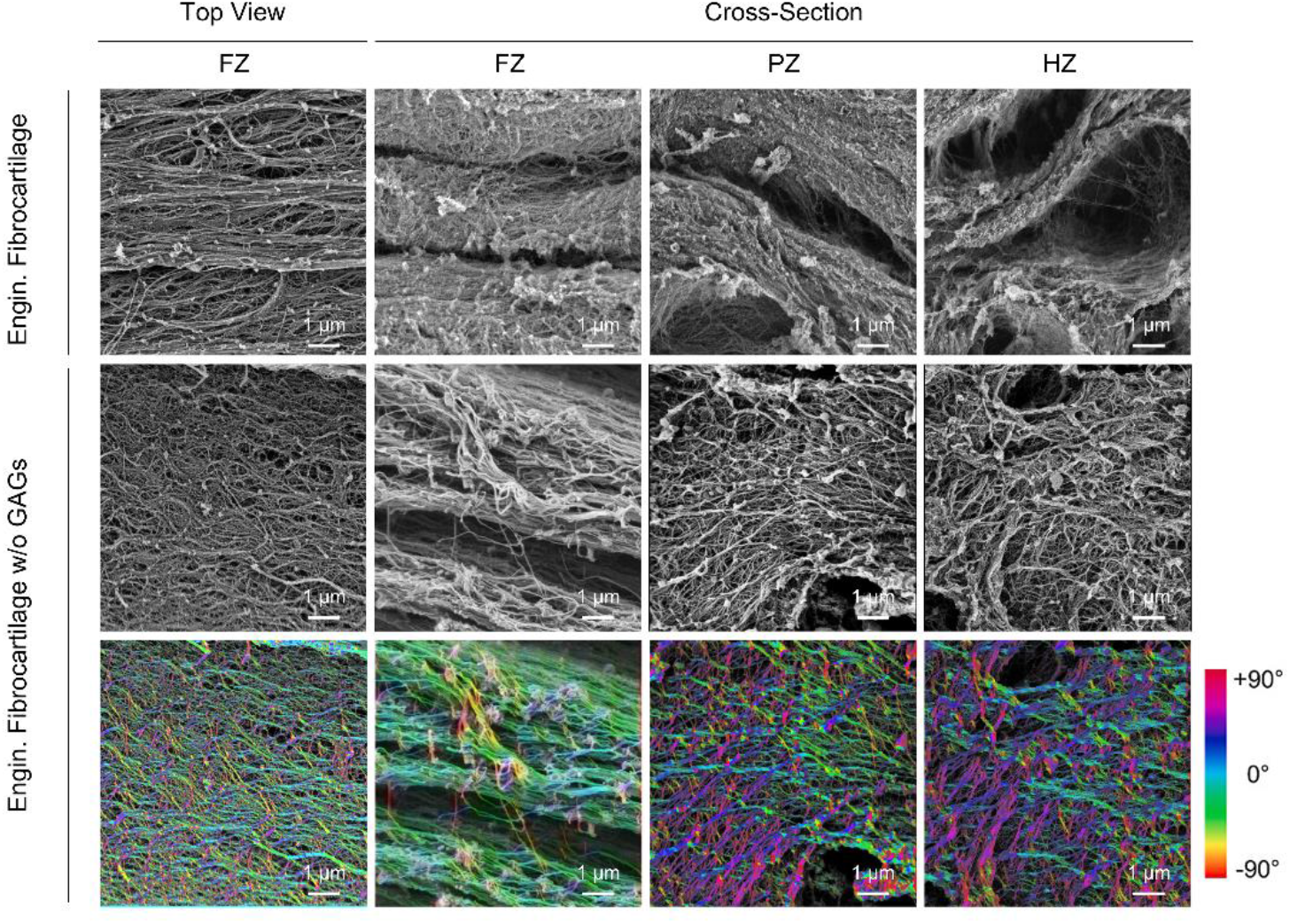
High-magnification scanning electron microscopy observations of collagen fibril organization in guided engineered tissue. Collagen fibrils were observed in the fibrous zone (FZ), proliferative zone (PZ), and hyaline zone (HZ). Observations were made in untreated samples (top row) and after glycosaminoglycan removal (w/o GAGs) by serial enzymatic digestion (middle row). The latter observations were color-coded to indicate fibril orientation (bottom row).

### 8. Micromechanical Properties of Engineered and Native Tissues

The micromechanical properties were examined in a depth-dependent manner in both guided engineered and native tissues (**Figure 8-A**). Young’s modulus, energy dissipation, and adhesion force were extracted from the retraction curve (**Figure 8-B**). Engineered tissues showed statistically significant depth-dependent mechanical properties, with Young’s modulus increasing from the surface to depth (**Figure 8-C**). Conversely, the adhesion force (**Figure 8-D**) and energy dissipation (**Figure 8-E**) decreased from the surface to depth. Native tissues demonstrated an overall opposite trend, although these differences were not statistically significant (**Figure 8-F, G, and H**). No significant differences were found when comparing the Young’s modulus (**Figure 8-I**), adhesion force (**Figure 8-J**), and energy dissipation (**Figure 8-K**) of PZ and HZ between engineered (6.6 ± 4.4 and 10.4 ± 5.4 KPa) and native tissues (8.8 ± 4.4 and 10.2 ± 5 KPa). However, the Young’s modulus of the FZ was significantly lower in engineered tissue (4.2 ± 1.6 KPa) than in native tissue (13 ± 6.8 KPa). Conversely, both adhesion force and energy dissipation of the FZ in engineered tissue (4.0 ± 2.3 nN and 7.0 ± 3.8 fJ) showed statistically significant higher values compared to native tissue (0.96 ± 0.58 nN and 1.6 ± 0.94 fJ). Similar results were obtained from the approach curve and by processing the retraction curve while accounting for adhesion (**Supplemental Figure 12**) (**Supplemental Table 3**).

**Figure 8.**
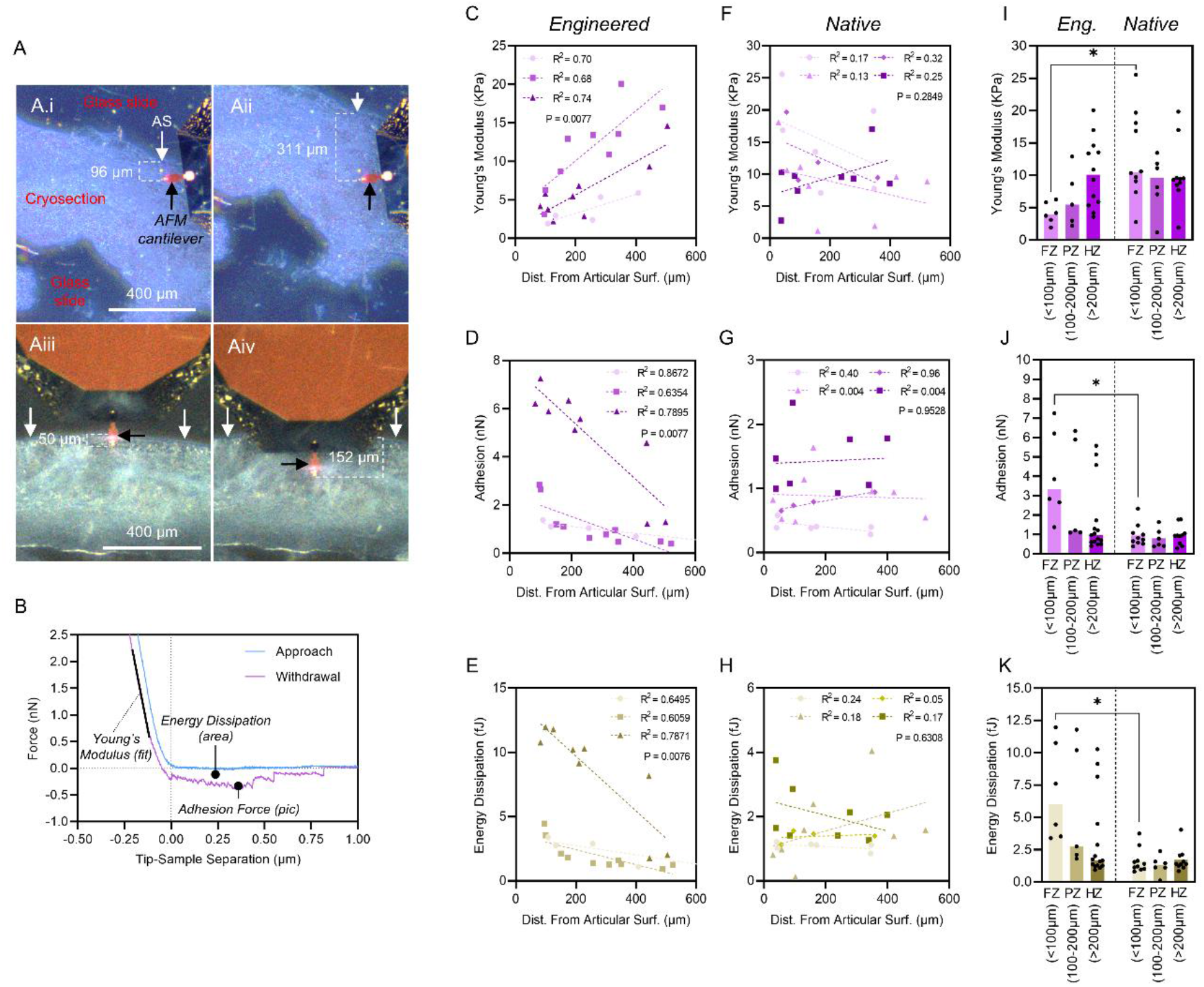
Micromechanical properties of guided engineered and native human mandibular condyle fibrocartilage. (A) Atomic force microscopy (AFM) was performed on (Ai, Aii) guided engineered and (Aiii, Aiv) native fibrocartilage cryosections in a depth-dependent manner to determine zonal micromechanical properties. White arrows indicate the articular surface (AS), and black arrows highlight the AFM cantilever. (B) Representative force-distance AFM curve obtained from guided engineered tissue, highlighting output values of interest. (C, F) Young’s modulus, (D, G) adhesion force from retraction curves, and (E, H) energy dissipation determined in a depth-dependent manner for (C, D, E) guided engineered and (F, G, H) native fibrocartilage tissues. Measurements were conducted on three and four distinct cryosections of the engineered and native tissues, respectively. (C-H) The results are represented as a trend bar for each cryosection, with multiple zones tested per cryosection depicted as data points. The P-values from the Pearson correlation test and the coefficient of determination (R^2^) are provided. The results in panels (I, J, and K) are reported as zone-specific values, with data displayed as bars indicating the median and each data point representing a measurement. Statistical significance is denoted by * for P < 0.05 (Kruskal-Wallis test followed by Dunn’s multiple comparisons test).

## Discussion

Although the knee joint and TMJ experience a similar incidence of cartilage disorders, higher research funding, academic publications, and research translation have been reported in the knee orthopedics field (e.g. up to nine-fold higher grant counts and six-fold higher article counts) [4]. This lack of primary research in TMJ tissue engineering has led to a shortage of funding and new therapeutics, creating a vicious cycle [24]. However, by using knee tissue engineering as a template, various approaches and suggestions could promote the development and implementation of tissue engineering-based therapeutics for the TMJ [4].

DWE is mostly known as melt-electrowriting (MEW) which involves melting the polymer before extrusion [25]. Biomimetic hyaline articular cartilage tissues have been produced by guiding the growth of spheroids within MEW scaffolds [18]. Additionally, by altering the aspect ratio of the porosities, biomimetic knee meniscus tissues have also been engineered [26]. Both approaches relied on spheroid self-assembly within the scaffolds. Given the structure of the TMJ condylar cartilage ECM, a rational adaptation of this biofabrication strategy was to seed preformed spheroids into the scaffolds to mimic the random organization of the PZ and HZ. This strategy was first applied by McMaster *et al*. for adipose tissue engineering, where they introduced "catching fibers" in MEW scaffolds to retain individual AD-MSC spheroids per pore [21]. Here, the same principle was applied to retain multiple AD-MSC spheroids in each pore of the dual-network structure. The structure was produced using solvent-based DWE instead of MEW, allowing the generation of millimeter-sized arrays of low-diameter polymeric fibers. Interestingly, the diameter of the catching fibers (≈ 14 µm) was larger than that of the main mesh (≈ 5 µm). This difference likely resulted from the spreading of the initial fiber layers onto the silicon wafer, followed by rapid rounding due to polymer-polymer repulsion during stacking.

Traditionally*, in vitro* cartilage tissue formation has been achieved using spheroids with a high cell count (≥ 2 x 10^5^ cells) [27,28]. Recent studies have significantly reduced this number (> 1,000 cells) [22,29] to align with the diffusion scale of signals during early developmental events such as condensation (≈ 80 - 200 µm) [30]. To balance these factors with production capabilities, we used spheroids containing approximately 3,000 cells with a diameter of approximately 300 µm to generate *in vitro* fibrocartilage. These spheroids were found to be homogeneous in ECM deposition and cell morphology, whereas larger spheroids containing 6,000 cells exhibited ECM depletion at the core and hypertrophic cell morphologies. Spheroids were assembled only 48 hours after condensation, as longer condensation periods before fusion led to inhomogeneous tissue fusion, evidenced by tenascin deposition between AD-MSC spheroids [27]. However, the use of early mature spheroids is challenging, as their assembly results in complete fusion, thereby limiting control over tissue morphology [27]. The DWE scaffold provided control over assembly constriction, enabling the assembly of early mature AD-MSC spheroids. Such control has also been demonstrated through compression molding [27,28] and by scaffolding spheroids into structures produced by two-photon polymerization 3D printing [31].

The DWE scaffold further regulated tissue growth, with the catching fiber network imposing clear limitations at the bottom, whereas tissue overgrowth occurred on the top of the structure, which remained unconstrained. Consequently, a sizeable graft with a diameter of 10 mm and polymeric volume fraction of approximately 0.6 % could be produced. In relation to the mandibular condyle, this graft represents around 40 % of the articular surface area [23]. In this study, approximately 13 × 10⁶ cells were seeded per graft, and around 34 × 10⁶ cells would be needed for complete mandibular condyle resurfacing. Pak *et al*. reported that the number of nucleated cells per milliliter of fat tissue ranges from 5 × 10⁵ to 2 × 10⁶ cells, with 1 to 10 % being AD-MSCs [32]. Considering an averaged 260 ml fat tissue volume collected for cell therapy in an autologous context [33], this translates to approximately 13 to 52 × 10⁶ early-passaged AD-MSCs available for engineering TMJ fibrocartilage, aligning with the cell quantities used in this study and those estimated for condyle resurfacing. Interestingly, the use of minimally expanded AD-MSCs has recently been applied in the engineering of phalangeal grafts to reduce manufacturing time and preserve chondrogenic potential [34].

Our results demonstrate that both adult native and engineered guided tissues exhibited a continuous FZ over the articular surface, composed of flattened cells and nuclei - a feature also described at the fetal stage in the human mandibular condyle [35]. Although the FZ in the engineered tissues was thinner, its thickness remained within the same order of magnitude as that of the native tissue. Interestingly, while only two layers of flat cells were observed in the articular surface of hyaline tissue grown from self-assembled spheroids in MEW scaffold [18], the engineered guided tissues in our study possessed a thicker FZ with multiple layers of flattened cells within the same culture period. This finding underscores the role of the timing of aggregate formation in the engineered tissue structure. In contrast, the engineered tissues displayed an abnormally thick HZ compared with the native tissue. The thickness of fibrocartilage on the mandibular condyle is known to vary anatomically [36] and tends to decrease with age [37]. Hence, histomorphometric measurements made by others showed a combined thickness of the PZ and HZ in human condylar fibrocartilage of approximately 600 μm [23], which aligns with the measurements observed in the guided engineered tissues. In the native tissue, the PZ was distinctly identified by a higher cell density, whereas the engineered tissues showed comparable cell densities between the PZ and HZ. Despite these differences, similar proliferative capacities were observed between engineered and native tissues.

Guided tissues cultured for 28 days exhibited GAG content comparable to that of native tissues from adult human donors. Spatial deposition of GAG, as observed by alcian blue staining, was biomimetic to that found in native human tissue but also rat TMJ [38]. Guided engineered tissues displayed a fibrocartilaginous composition and stained positively for both type I and type II collagens. However, type I collagen, the primary component of fibrocartilage, was distributed throughout the tissue section, while type II collagen was predominantly expressed in the HZ, similar to adult human tissue [37]. Other major ECM components, aggrecan core protein and fibronectin, also exhibited similar deposition patterns between the guided engineered and native tissues. Nevertheless, the stronger immunofluorescence detected in native tissue for type I collagen and aggrecan core protein indicated a lack of ECM content in engineered tissues, further evidenced by the 10-fold lower total collagen content in engineered tissues than in native tissue. Despite the lower total collagen content after 4 weeks of culture, the collagen network organization mimicked that observed in human TMJ condylar fibrocartilage. Specifically, tissue growth resulted in a randomly organized network in the PZ and HZ, which transitioned abruptly to strongly aligned layers of collagen fibrils running parallel to the surface in the FZ [23]. Again, tissue resulting from the growth of in-situ assembled spheroids exhibited a completely different collagen architecture, akin to a Benninghoff-like collagen alignment [18], further highlighting the impact of spheroid formation timing on tissue structure. Achieving native-level of collagen could be facilitated by administering exogenous lysyl oxidase (LOX), an enzyme involved in collagen cross-linking, which has been shown to improve the tensile properties of engineered cartilage [39]. Additionally, combining LOX with chondroitinase-ABC and TGF-β1 in a sequential treatment regime increased collagen content and enhanced *in vivo* integration of engineered fibrocartilage tissues [40]. Extending the culture period [41] and priming with bone morphogenetic protein (BMP) 6 [42] could also further enhance the overall ECM synthesis.

Bulk mechanical testing of similar tissue composites showed that polymeric fibers reinforced tissue under tension, while newly synthesized ECM was responsible for the tissue’s compressive properties [18]. In the present study, we adopted a different approach by testing the micromechanical properties of fibrocartilage and comparing them to those of the native tissue. In the latter, Rang *et al*. reported a higher elastic modulus in the FZ than in the HZ across five species, except in humans, where 35 kPa and 61 kPa were measured in the FZ and HZ respectively. This was investigated in human specimens obtained from patients undergoing joint replacement surgery with potential TMJ dysfunction, which might have influenced the fibrocartilage properties, as in the present study. Nevertheless, the overall trend here aligned with findings across other species, suggesting that healthy regions were tested, with values of the same order of magnitude as those reported by Rang *et al*.

Histological investigations did not show a significant difference in the overall collagen content across zones in native tissues; however, type I collagen immunostaining and PLM demonstrated a clear trend toward higher content and fibril alignment in the FZ. This trend is consistent with the AFM results, which showed a higher, though not statistically significant, elastic modulus in the FZ than in the PZ and HZ. In engineered guided tissues, significant zonal differences were found in the overall collagen content, but these did not match the trend in elastic modulus observed *via* AFM. Nevertheless, type I collagen immunostaining indicated a trend toward a lower content in the FZ. Notably, the organization of the collagen network itself may intrinsically influence AFM measurements, since the parallel alignment of fibrils in the FZ may reduce cantilever adhesion, leading to a lower modulus, whereas the anisotropic arrangement in the PZ and HZ, with a larger contact surface area, could naturally produce the opposite effect.

We found that the primary difference in micromechanical properties between engineered and native fibrocartilage was the two-fold lower elastic modulus in the FZ of engineered tissues, supported by the higher total collagen and type I collagen immunostaining in native tissue. This finding suggests that enhancing the engineered graft should focus on improving the maturity of the FZ, for instance by exposing the developing tissue to joint-mimicking motions. Indeed, a loading regime applied by a humanoid robot shoulder joint was shown to influence the gene expression profile of human fibroblasts grown on fibrous scaffolds [43]. Furthermore, cyclic loading affected cell-driven collagen organization and maturation in fibroblasts derived from the anterior cruciate ligament [44]. Given the complexity of potential mechanical load combinations and their influence on tissue maturation, Ladner *et al*. used a full factorial design of experiments to investigate the effects of combined shear and compressive strains provided by cylindrical or spherical counterfaces on biological markers [44]. This type of modular bioreactor and screening strategy could help optimize loading protocols for FZ maturation.

The implementation of controlled spheroid deposition is also a crucial aspect of this biofabrication strategy. To date, various technologies have been described [45–48] and implemented for engineering osteochondral grafts [49]. From a tissue engineering perspective, controlled deposition could help reduce the HZ thickness to an average physiological level. Furthermore, anatomical studies have shown that fibrocartilage exhibits varying thickness depending on its location on the mandibular condyle [36]. Combined with the controlled variation in DWE scaffold thickness, this approach could also accommodate anatomical tissue variations. To facilitate implantation and integration, the tissue graft could be combined with harder structures, such as metals or ceramics. While generating the DWE scaffold on a complex curved implant seems feasible [50,51], followed by spheroid deposition, an unconventional approach like hard–soft electroadhesion [52] could also be explored considering that cells have been shown to survive when extruded through an electric field [53].

## Conclusion

Building on recent advances in engineering biomimetic hyaline cartilage, we demonstrated that combining early-mature human AD-MSC spheroids with polymeric scaffolds resulted in the formation of biomimetic TMJ fibrocartilage assembloids. Within just 4 weeks, these tissue analogs exhibited native-like spatial ECM deposition and collagen network organization, as well as similar micromechanical properties. From a fundamental standpoint, these results highlight that modifying the timing of cell aggregate integration within the scaffold is another key factor in reorienting ECM, alongside scaffold architecture, further showcasing the versatility of this biofabrication strategy in achieving “orthopaedic tissues by design.” From a translational perspective, the protocol developed here offers a simple and efficient *in vitro* culture system for priming fibrocartilage grafts using a clinically relevant number of cells within a few weeks. This can revolutionize the mitigation of current clinical challenges such as TMJ reconstruction.

## Experimental Section

### 1. Cell Isolation, Expansion, and Spheroid Production

Human AD-MSCs isolated from abdominal surgical liposuction (female, 44 years old, body mass index 18.51 kg/m2) were purchased from the cell and tissue bank of the *Hospices Civils de Lyon* (Lyon, France) according to French regulations, with a declaration to the French Minister of Research (No. AC 2008-162 and DC No. 2019-3476). Cells were seeded at 2.5 x 10^3^ cells/cm^2^ and cultured up to passage five in Mesenchymal Stem Cell Growth Medium 2 (MSC-GM2) (Promocell, #C-28009).

Silicone inserts with microwells were fabricated in-house for spheroid generation. Initially, a positive mold accommodating 401 microwells of 0.4 mm diameter and 0.8 mm height **(Supplemental STL 1)** and a casting frame **(Supplemental STL 2)** were 3D-printed using VeroClear material with a PolyJet inkjet printer (Stratasys, USA). Silicone (Elkhem Silicone, #RTV-141) was poured over the positive mold placed in the casting frame and crosslinked at 60 °C for 4 h. Following additional curing at 150 °C for 30 min and steam sterilization, the inserts were placed in a 6-well plate and centrifuged with an anti-adherence solution (StemCell Technologies, #07010) at 250 g for 5 min. Following a further 5- minute incubation, the inserts were washed with phosphate-buffered saline (PBS). Cells were added at a density of 1,500, 3,000, or 6,000 cells per microwell with 4 ml of chondrogenic medium per insert and centrifuged at 250 g for 5 min. Chondrogenic medium consisted of high glucose DMEM (Gibco™, #10569010) supplemented with 1 X insulin-transferrin-selenium (Gibco™, #41400045), penicillin (100 U/mL) and streptomycin (100 μg/ml), 40 μg/ml L-proline (#P5607), 50 μg/ml L-ascorbic acid-2- phosphate (#A8960), 4.7 μg/ml linoleic acid (#L5900), 1.5 mg/ml bovine serum albumin (BSA, #A7906), 100 nM dexamethasone (#D8893) (all from Sigma-Aldrich), and 10 ng/ml human transforming growth factor-beta (TGF-b) 3 (PeproTech, #100-36E). Spheroids were allowed to self-assemble for 48 h without medium changes before harvest.

### 2. Direct-Writing Electrospinning (DWE)

Scaffolds were fabricated using a customized DWE printer [54] within a controlled environment (20 ± 1 °C and 40 ± 1 % humidity). A 32 % (w/v) solution of medical-grade PCL (Corbion, #PC12) dissolved in 1,1,1,3,3,3-hexafluoro-2-propanol (CAS #920-66-1) was used as the material. The solubilized PCL was extruded at a rate of 0.02 ml/h from a fixed 2 ml syringe equipped with a 21G needle onto a CZ silicon wafer (Siltronix) secured on the collector plate. A voltage of 1.8 KV was applied between the nozzle and the collector plate, which was positioned at a distance of 3 mm. The collector plate moved at a speed of 550 mm/s in both the X and Y directions. Additionally, the collector plate was incrementally lowered by 30 µm every 50 layers.

Before completing the 3D structure, the stability of the PCL jet was ensured by printing 20 lines, which were examined for deviations in the fiber diameter or pulsing. The scaffolds consisted of two types of meshes: a box-like structure measuring 19 × 19 mm, termed the main mesh, with each fibrous layer oriented at 90° to the previous layer with 0.9 mm spacing. Additionally, a second mesh measuring 23 × 23 mm was included, with each fibrous layer also oriented at 90° to the previous layer, but with a spacing of 0.15 mm between them. The scaffold design comprised five layers of the main mesh, followed by three layers of the second mesh at 45°, and concluded with 295 layers of the main mesh, resulting in a total height of 1 mm.

The scaffolds were secured for cell culture using custom 3D-printed inserts. Initially, a two-part inner insert with an inner diameter of 6 mm was 3D-printed with FibreTuff® using a Prusa Mini printer (Prusa Research Technology, Czech Republic). Additionally, an outer insert designed to fit a 6-well plate was 3D-printed with Makrolon® polycarbonate using Freeformer technology (Arburg, Germany) (**Supplemental Figure 13**). Both materials were subjected to steam sterilization. DWE scaffolds were immersed in 70 % ethanol overnight, mounted between the two parts of the inner insert, and placed in the outer insert. The entire assembly was transferred to a sterile 6-well plate cell culture dish. In this setup, the DWE scaffold was under tension and was elevated 1 mm above the bottom of the culture plate. Finally, the setup was subjected to ozone sterilization for one hour with 30 min of ultraviolet exposure in parallel.

### 3. Construction of Assembloids

The number of spheroids seeded for each construct was determined with a previously established equation [22], considering the spheroid volume and the volume of the final construct. Spheroids with a diameter of approximately 300 µm (3,000 cells/spheroid) were seeded into constructs measuring 10 mm in diameter and 1 mm in height. This resulted in 4,468 spheroids/structures, with a density of approximately 57 spheroids/mm^3^. After 48 h of condensation, spheroids resuspended in chondrogenic medium were manually seeded into DWE scaffolds or a silicone mold of equivalent volume and geometry, pre-treated with an anti-cell adhesive coating as described above. The developing fibrocartilage was cultured in 10 ml chondrogenic medium under physioxic conditions for 4 weeks, with full media exchange performed every 2-3 days. Following *in vitro* maturation, the gross morphology of the tissues was imaged using a Nikon SMZ1270 stereomicroscope equipped with a DS- Fi3 camera.

### 4. Native Human Mandibular Condyle

Human TMJ samples were collected from a 40-year-old female patient who underwent TMJ replacement. The collection was conducted following written informed consent from the donor and in full compliance with local ethics guidelines as well as national and European Union legislation concerning human sample collection, handling, and personal data protection (CODECOH N°DC-2021- 4470).

The TMJ sample was decalcified (Osteomoll, Sigma-Aldrich, #101736) before histological processing. Type IV collagen immunostaining was performed on human mandibular condyle tissue sections to identify regions of interest. Although type IV collagen has been found in the pericellular matrix of chondrocytes in rat TMJ [55], it is also described as a characteristic feature of the degenerative lesion in human TMJ [37]. Positive staining was observed in the subchondral bone beneath the condylar cartilage, lining the blood vessels. No staining was observed in the condylar cartilage, except in zones where the articular surface was disrupted, indicating a degenerative lesion (**Supplemental Figure 5**). Consequently, further analysis and measurements were confined to disease-free regions that lacked positive type IV collagen staining and appeared histologically similar to healthy tissues from young adults [37].

### 5. Biochemical Analyses

Engineered and native fibrocartilage was weighed and digested using papain (3.88 units/ml, Sigma-Aldrich, #P3125) in a buffer containing 100 mM sodium phosphate, 5 mM ethylenediaminetetraacetic acid (EDTA, pH 6.5), and 10 mM L-cysteine hydrochloride (Sigma-Aldrich, #C7880) at 60 °C with agitation. DNA content was quantified using the Hoechst Bisbenzimide 33,258 dye assay (Sigma-Aldrich, #DNAQF). GAG content was measured using a dimethylmethylene blue dye binding (DMMB) assay (Biocolor Ltd, Blyscan™). The total collagen content was estimated by measuring hydroxyproline levels and applying a hydroxyproline-to-collagen ratio of 1:7.69. This involved the oxidation of hydroxyproline residues (OHP) using chloramine-T (Sigma-Aldrich, #857319), followed by the formation of a chromophore with dimethylaminobenzaldehyde (Sigma-Aldrich, #109762) [56].

### 6. Histological and Immunohistochemical Analyses

Tissues were rinsed in 1× PBS and fixed in paraformaldehyde (Antigenfix solution, DiaPath, #P0016) overnight at 4 °C. After rinsing in PBS, the samples were dehydrated in a graded series of ethanol from 30 to 70 % (v/v) and stored at 4 °C until paraffin embedding, followed by sectioning at 5 μm. The sections were stained with H&E to study tissue and cell morphologies, alcian blue & nuclear fast red to reveal the presence of GAGs and cells, and picrosirius red to visualize the collagen content.

Collagen types I, II, and IV, along with aggrecan, fibronectin, and Ki-67, were assessed using standard immunostaining protocols. The antigen retrieval methods and the primary and secondary antibodies used for immunostaining are listed in **Table 2**. Briefly, rehydrated tissue sections were treated with either hyaluronidase (800 U/mL, Sigma-Aldrich, #R3506), pepsin (Sigma-Aldrich, #R2283), trypsin (GBI Labs, #E05-18), or Tris-based buffer (#CC1, Roche Diagnostic). Subsequently, the sections were incubated with BlockAID (Invitrogen, #B10710) for 20 min to block non-specific binding sites. The tissue sections were then incubated overnight at 4 °C in a humidified chamber with the primary antibody diluted in the blocking solution. Afterward, they were treated for 1 h at room temperature with the secondary antibody diluted in blocking solution. Finally, the nuclei were stained with 300 nM 4’,6-diamidino-2-phenylindole dilactate (DAPI) (Invitrogen, #D3571) for 10 min at room temperature, and the slides were mounted using Fluoromount (Sigma-Aldrich, #SLCQ9183).

**Table 2.** Antigen retrieval method, primary and secondary antibodies used for immunostaining.

|  | Target Protein | Antigen Retrieval Method | Antibody Properties | Dilution | Reference |
| --- | --- | --- | --- | --- | --- |
| Primary Antibodies | Type I collagen | 1. Hyaluronidase 800 U/ml, 37 °C, 30 min<br>2. Pepsine, RT, 15 min | Rabbit polyclonal anti-type I collagen | 1:250 | Novotec (#20111) |
|  | Type II collagen | Pepsine, 37 °C, 30 min | Rabbit polyclonal anti-type II collagen | 1:50 | Novotec (#20211) |
|  | Aggrecan | Hyaluronidase 800 U/ml, 37 °C, 30 min | Rabbit polyclonal anti-aggrecan core protein | 1:200 | Novotec (#CSPG 10.04.14) |
|  | Fibronectin | Trypsin, 17min, 37 °C | Rabbit polyclonal anti-fibronectin | 1:150 | Novotec (#24911) |
|  | Type IV collagen | 1. Hyaluronidase 800 U/ml, 37 °C, 30 min<br>2. Pepsine, RT, 15 min | Rabbit monoclonal anti-type IV collagen | 1:150 | Abcam (#ab214417) |
|  | Ki-67 | Tris based buffer | Rabbit monoclonal anti-Ki-67 | 1 :1000 | Abcam (#ab16667) |
| Secondary Antibodies |  |  | Alexa Fluor 488-conjugated anti-Rabbit | 1:500 | Invitrogen (#A11008) |
|  |  |  | Anti-Rabbit IgG | NA | Vector Laboratories (#MP-7601) |

### 7. Polarized Light Microscopy (PLM)

Rehydrated tissue sections were incubated at 37 °C for 18 hours with 1000 U/mL bovine testicular hyaluronidase (Sigma-Aldrich, #H3506) prepared in 0.1 M phosphate buffer pH 6.9 to remove proteoglycans so birefringence was only caused by collagen fibrils [57,58]. Sections were then stained with 0.1 % (w/v) picrosirius red, mounted with DPX (all from Sigma-Aldrich), and imaged using a Zeiss Axioscan 7 slide scanner.

### 8. Scanning Electron Microscopy

For high-level observation, samples were fixed in 3 % (v/v) glutaraldehyde in 0.1 M cacodylate buffer at 4 °C for at least 12 h. After rinsing twice in 0.1 M cacodylate buffer for 10 min, samples were dehydrated in a series of graded ethanol baths, immersed twice in hexamethyldisilazane (Sigma-Aldrich, #440191) for 30 min, and allowed to dry completely overnight before imaging. The samples were imaged using a TM4000II tabletop SEM (Hitachi High-Tech, Japan). Serial enzymatic digestion was used to remove GAGs for high-resolution visualization of the collagen network, providing an unobstructed view of the collagen fibrils [59]. The samples were imaged using a Merlin Compact VP microscope (Zeiss, Germany).

### 9. Image Analysis

All histological sections, including the PLM, were imaged with an Axioscan 7 slide scanner (Zeiss, Germany). Image analyses were performed using QuPath [60] and ImageJ [61] softwares. The size of spheroids was determined by measuring the area, then calculating the diameter of a circle with an equal projection area [62]. For histological staining quantification, the Color Deconvolution plugin in ImageJ was used with built-in settings to extract the blue channel from alcian blue and nuclear fast red images [63]. The zonal mean gray value was then measured in the processed alcian blue and nuclear fast red images, as well as in the picrosirius red-stained tissue sections. The average orientation, dispersion, and coherency of collagen fibrils in the PLM images were assessed using the OrientationJ plugins in ImageJ. Color orientation mapping for PLM and SEM pictures was obtained with Directionality plugin in ImageJ [64]. Quantification of the DAPI-stained nuclei was performed using the Particle Analysis function in ImageJ. Quantification of Ki-67-positive nuclei was performed using QuPath software. The immunofluorescence intensity of type I and type II collagens, aggrecan core protein, and fibronectin was determined using ImageJ by subtracting the product of the area of the selected zone and the mean fluorescence of background readings from the integrated density. False-color SEM images were obtained using Adobe Photoshop (Adobe Inc., USA).

### 10. Atomic Force Microscopy (AFM)

After 4 weeks of chondrogenic induction, excess medium was removed, and samples were stored at - 80 °C without prior chemical fixation. After thawing and removal of excess media, the samples were successively immersed in 15 % and 30 % (w/v) sucrose solutions at 4 °C overnight. Subsequently, the samples were immersed in a 50:50 (v/v) mixture of 30 % sucrose and Optimal Cutting Temperature (OCT) compound at room temperature for 2 h. Following mounting, the samples were frozen at -20 °C for at least overnight. Cryosections of 50 µm thickness were deposited on Epredia™ SuperFrost Plus^©^ Gold glass slides (Thermo Scientific, #K5800AMNZ72), left to dry at room temperature overnight, and then stored at -20 °C. AFM measurements were performed using a JPK NanoWizard® 3 (Bruker, Germany) with a 10 µm diameter spherical probe (Bruker, #SAA-SPH-5UM) on tissue sections hydrated with PBS.

The calibration process comprised measuring the deflection sensitivity on a Saphir sample and configuring the spring constant of the cantilever as provided by the manufacturer. Force mapping was then conducted on tissue sections within a 50 x 50 µm area divided into 10 x 10 pixels. The approach was executed with a Z-length of 7.5 µm at an extension speed of 2 µm/s. The data were sampled at a rate of 2000 Hz with a set point of 10 nN. AFM curves were subsequently processed using JPK Data Processing software (JPK Instruments, Germany).

### 11. Statistical Analyses

Graphical representations and statistical analyses were performed using the GraphPad Prism software (version 9.00). Data representation and statistical tests are detailed in the figure legends. Nonparametric tests were used to compare groups with fewer than 10 data points. The data normality of groups with a larger number of data points was assessed prior to performing comparisons. Parametric tests were used when the data were normally distributed for all groups; otherwise, non-parametric tests were used. Statistical significance was set at a threshold of P < 0.05.

## Acknowledgements

The authors thank 3Deus Dynamics for providing access to TM4000II tabletop SEM. They also thank the *Centre d’Imagerie Quantitative Lyon-Est* (CIQLE) for helpful advice on histological processing. The authors are grateful to Marilyne Duffraisse from the *Institut de Génomique Fonctionnelle de Lyon* (IGFL) for arranging access to the Keyence VHX stereomicroscope. Special acknowledgment is given to Simone Bovio from the *Plateau Technique d’Imagerie/Microscopie* (PLATIM) of the *École Normale Supérieure* (ENS) *de Lyon* for his invaluable assistance with AFM experiment. The authors also thank the *Centre Technologique des Microstructures de l’Université Lyon 1* (CTµ) for their assistance with the Merlin Compact VP SEM. The authors thank Céline Thomann for her assistance with cell culture, and Jérôme Lafont for kindly providing the anti-human aggrecan core protein antibody. Finally, the authors are grateful to Louis Brochet, Edouard Lange, and Sanela Morand for arranging the collection of the human TMJ sample.

## Author contributions

Conceptualization: AD; Methodology: AD; Investigation: AD, LE, VS; Validation: AD; Formal analysis: AD; Data Curation: AD; Visualization: AD; Supervision: AD; Project administration: AD; Funding acquisition: CM; Writing - Original Draft: AD; Writing - Review & Editing: AD, LE, KB, VS, CM. The manuscript was approved by all authors.

## Declaration of interests

The authors declare that they have no known competing financial interests or personal relationships that could have appeared to influence the work reported in this paper.

## Notes

### Competing Interest Statement

The authors have declared no competing interest.

