## Supplemental for "Biofabrication of spatially organized temporo-mandibular fibrocartilage assembloids"

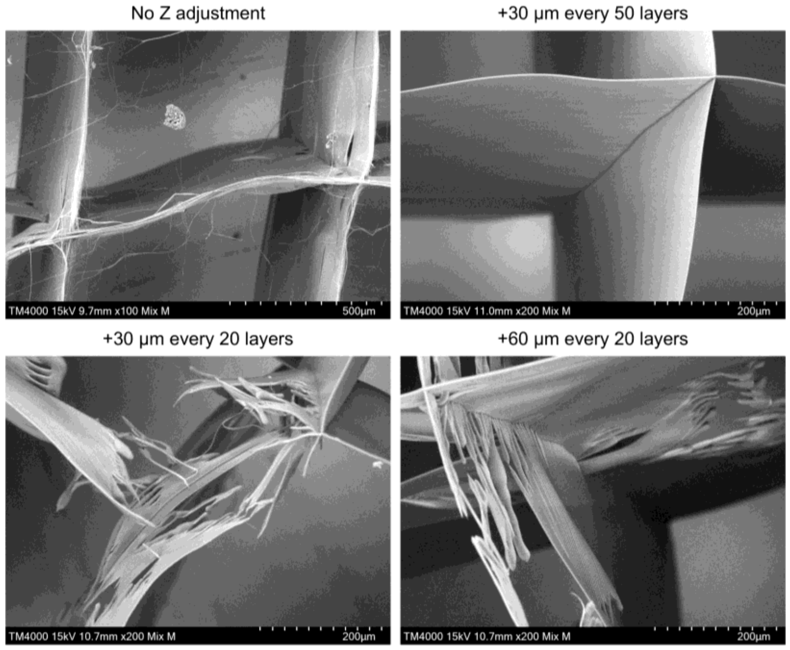


**Supplemental Figure 1**. **Effect of nozzle-to-wafer distance on the quality of fiber walls**. Maintaining a constant nozzle-to-wafer distance throughout the printing process resulted in pulsing and disruption of the jet. Increasing the nozzle-to-wafer distance by 30 µm every 50 printed layers produced consistent fiber walls. Higher distancing rates resulted in disrupted fiber walls.

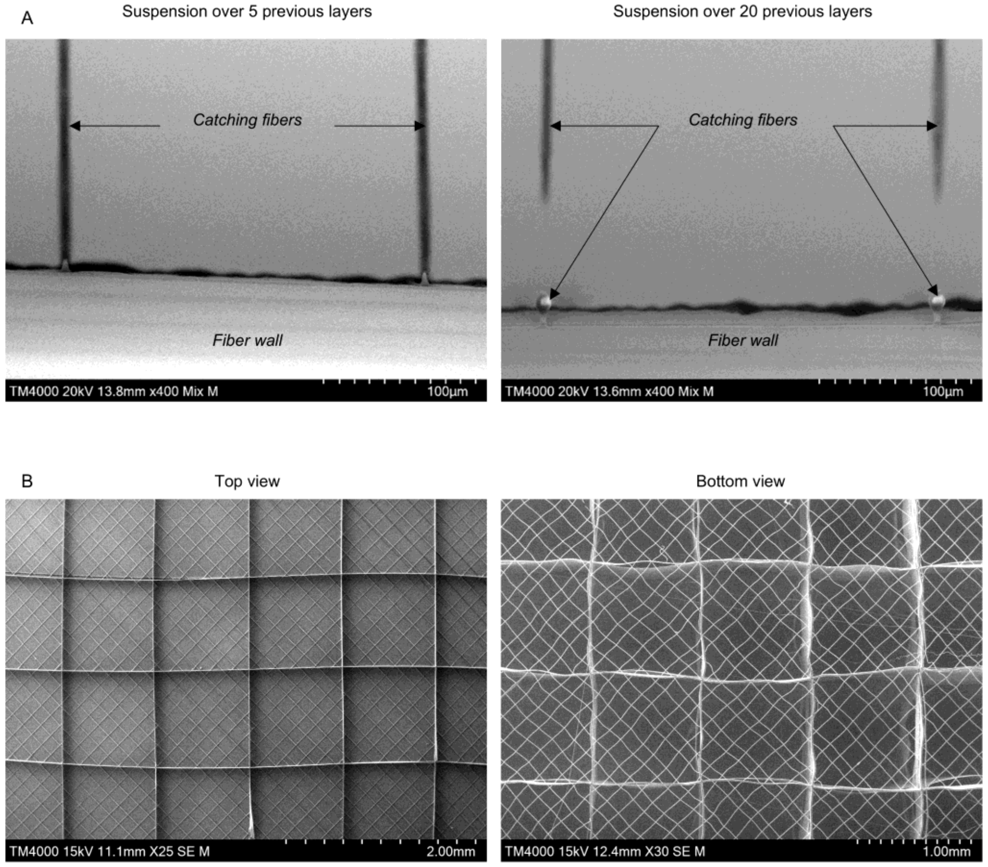


**Supplemental Figure 2. Scanning electron microscopy images of catching fibers and scaffolds.** (A) Quality of catching fibers as a function of the underlying printed layers. (B) Overview of printed scaffolds from the top and bottom perspectives.

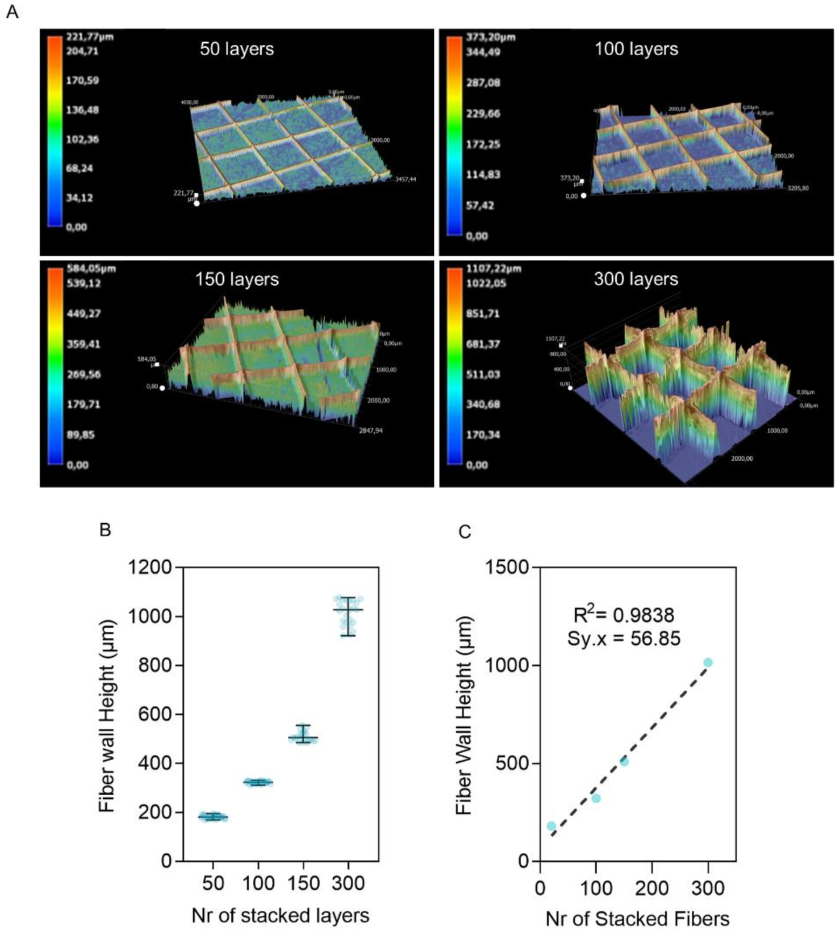


**Supplemental Figure 3. Measuring scaffold height with digital microscopy**. Scaffold height as a function of the number of printed layers was determined using a Keyence VHX 7000 digital microscope, enabling (A) 3D modeling of the structures and (B-C) quantitative evaluation of multiple fiber wall heights. (B) Graphical representation showing the mean of fiber wall height with the standard deviation. Each data point represents a fiber wall. (C) Simple linear regression analysis of the data set, with each data point representing the mean of fiber wall.

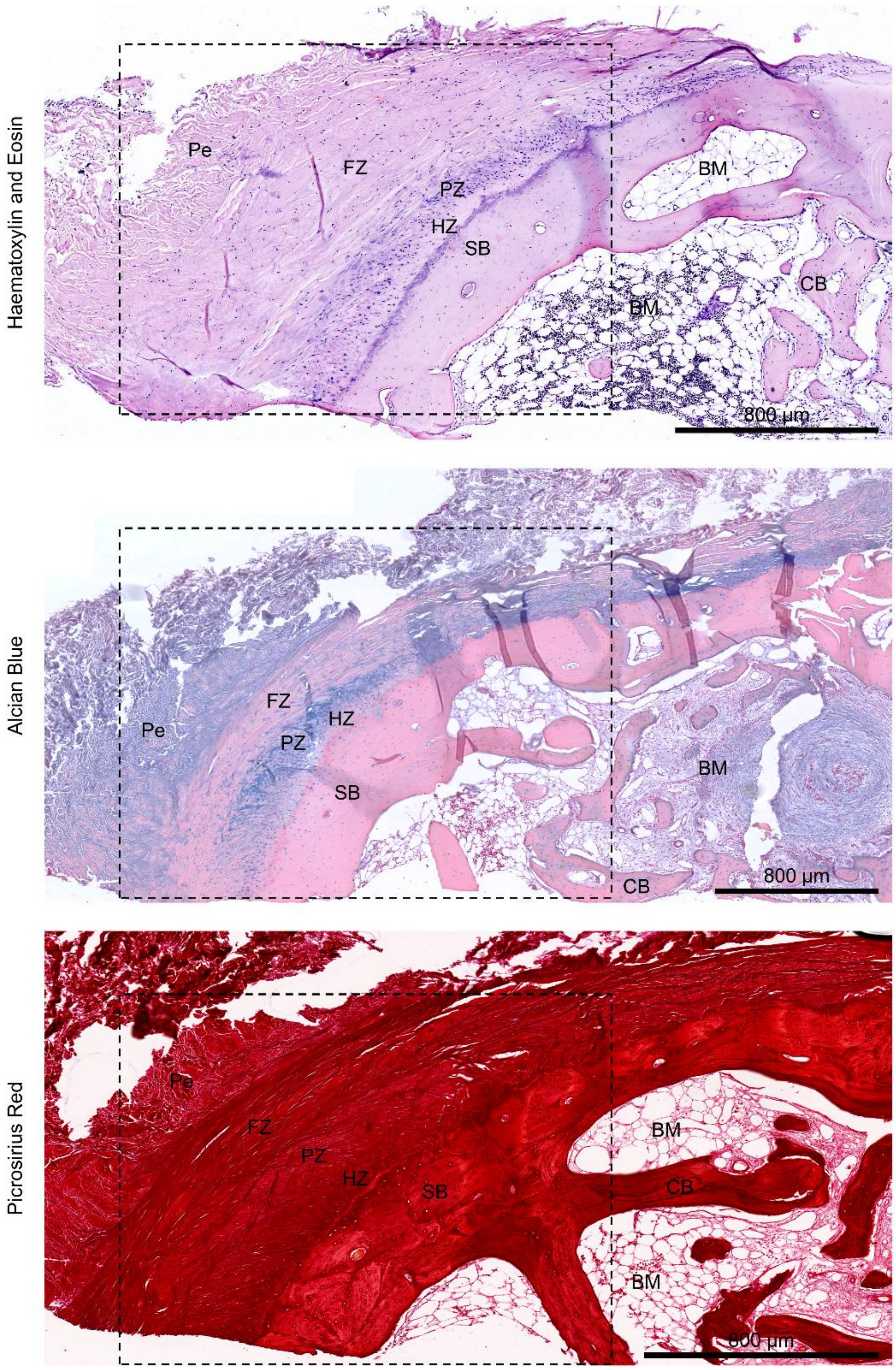


**Supplemental Figure 4. Large images of histologically stained human mandibular condyle**. Tissue was stained with (A) hematoxylin & eosin, (B) alcian blue & nuclear fast red, and (C) picrosirius red. Dotted lines show regions used for analysis and measurements. Pe, perichondrium; FZ, fibrous zone; PZ, proliferative zone; HZ, hyaline zone; SB, subchondral bone; BM, bone marrow; CB, cortical bone.

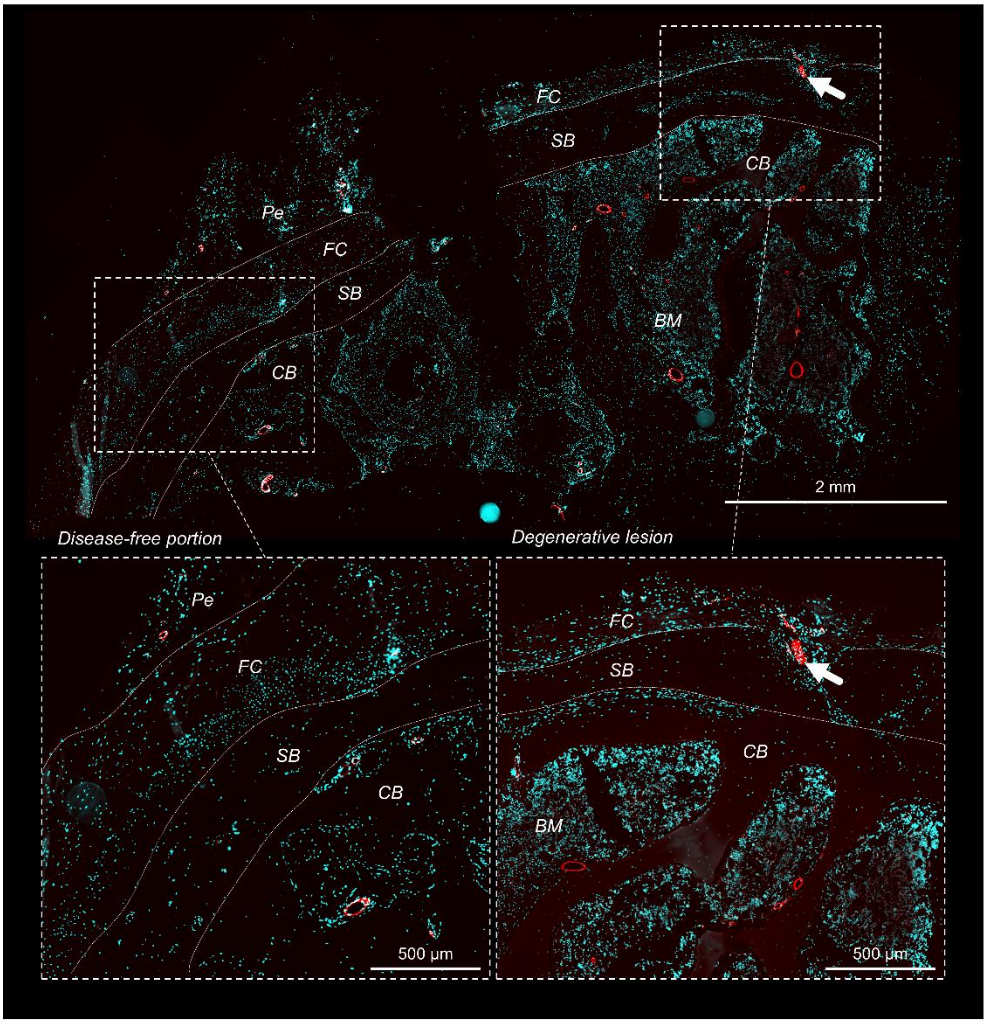


**Supplemental Figure 5. Immunofluorescence staining of the human mandibular condyle for type IV collagen.** Nuclei are indicated in cyan and type IV collagen-positive elements are shown in red. Since type IV collagen is a component of basement membranes in blood vessels, these structures are clearly visible in the cortical bone, the bone marrow and the perichondrium. No type IV collagen-positive staining is detected in disease-free portions of the fibrocartilage, while positive staining is indicated by a white arrow in degenerative lesions. Pe, perichondrium; FZ, fibrous zone; PZ, proliferative zone; HZ, hyaline zone; SB, subchondral bone; BM, bone marrow; CB, cortical bone.

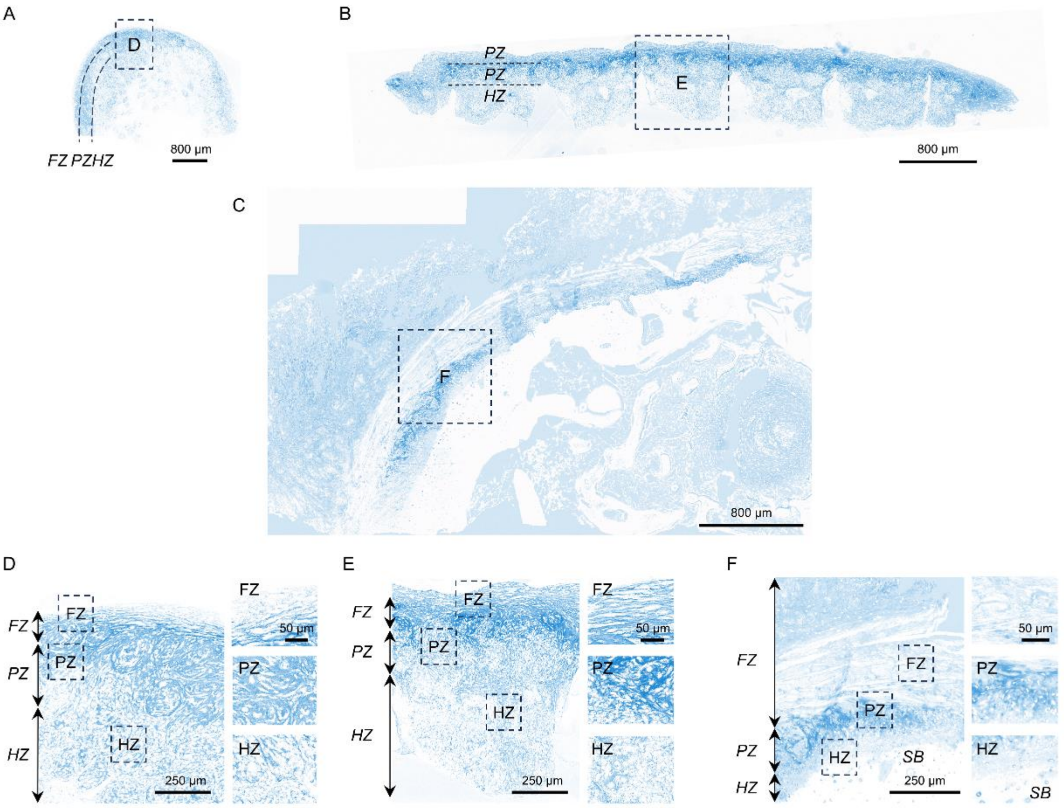


**Supplemental Figure 6. Alcian blue channel after color deconvolution of alcian blue and nuclear fast red histological images.** Image processing was performed using the dedicated plugin in FIJI with pre-determined parameters. (A-C) Low magnification images of (A) unguided and (B) guided engineered tissues, and (C) native human mandibular condyle. (D-F) High magnification images of (D) unguided and (E) guided engineered tissues, and (F) native human mandibular condyle fibrocartilage. FZ, fibrous zone; PZ, proliferative zone; HZ, hyaline zone.

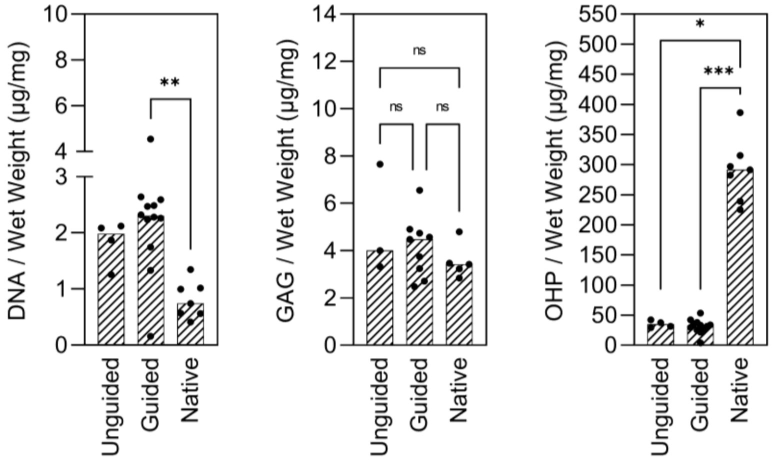


**Supplemental Figure 7**. **Biochemical properties of engineered and native fibrocartilage**. GAG, DNA, and total collagen (OHP) content normalized to wet weight. Each data point corresponds to a technical replicate with median shown as bars. Statistical significance was determined using Kruskal-Wallis test followed by Dunn’s multiple comparisons test. *P < 0.05, **P < 0.01 and ***P<0.001.

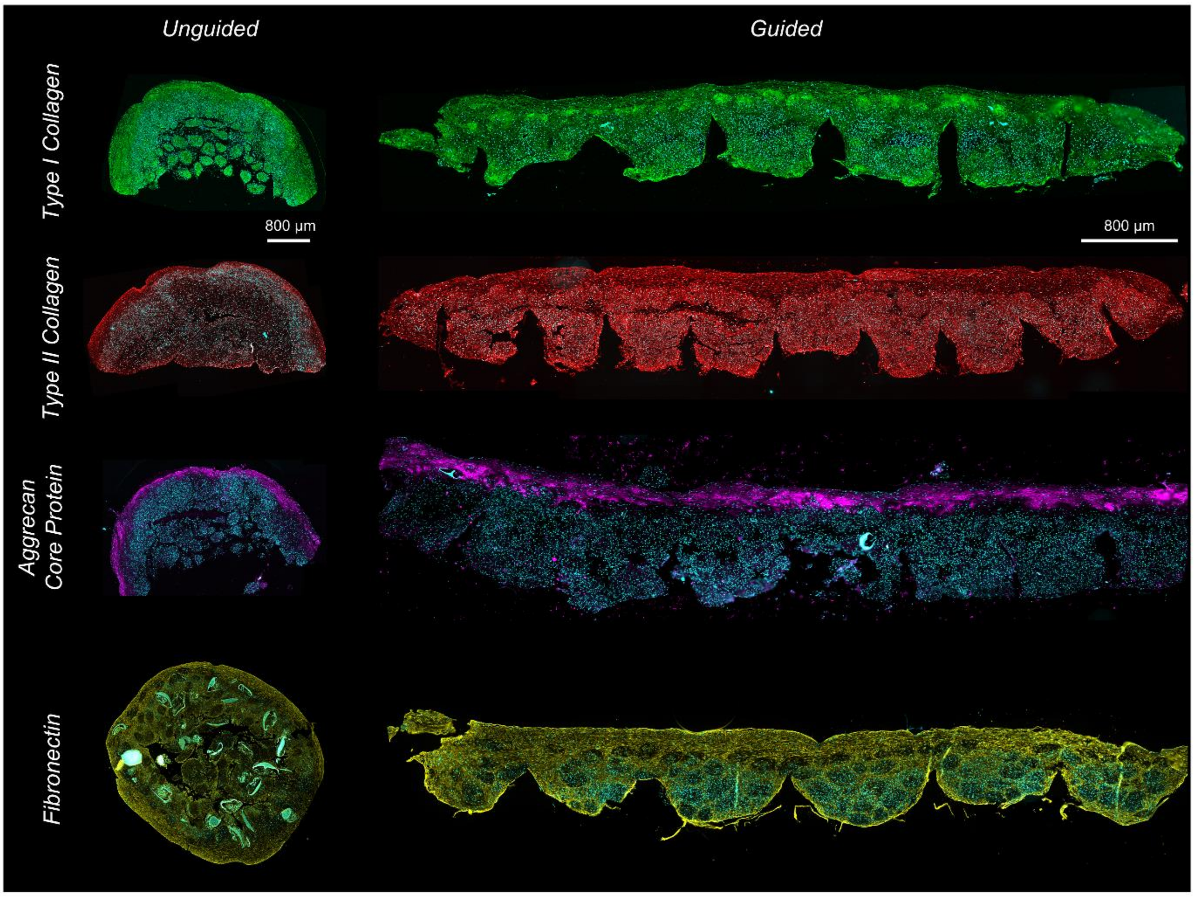


**Supplemental Figure 8. Large images of immunofluorescence staining of unguided and guided engineered tissues.** Nuclei are stained in cyan, type I collagen in green, type II collagen in red, aggrecan core protein in purple, and fibronectin in yellow.

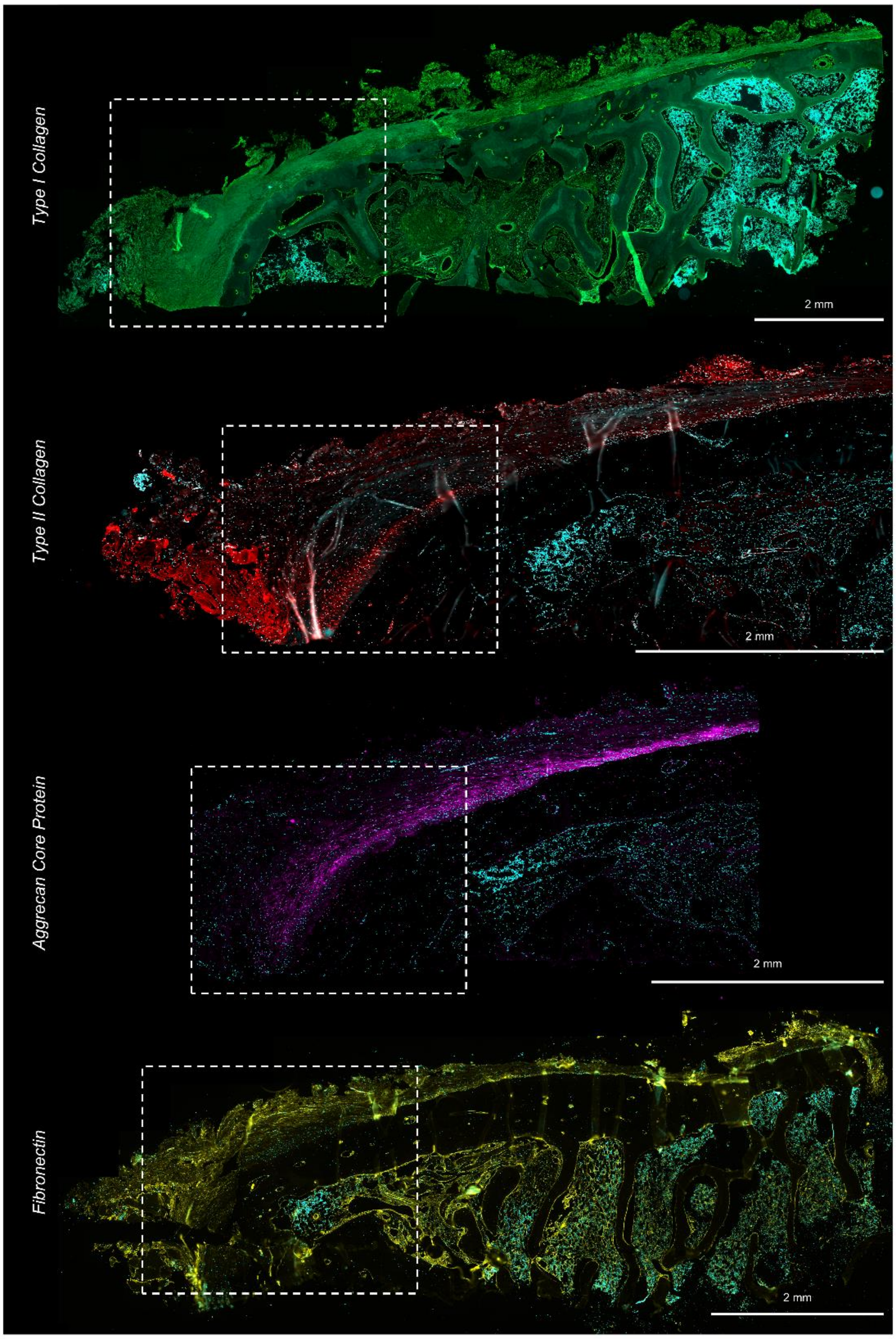


**Supplemental Figure 9. Large images of immunofluorescence staining of the human mandibular condyle.** Nuclei are stained in cyan, type I collagen in green, type II collagen in red, aggrecan core protein in purple, and fibronectin in yellow. Dotted lines show regions used for analysis and measurements.

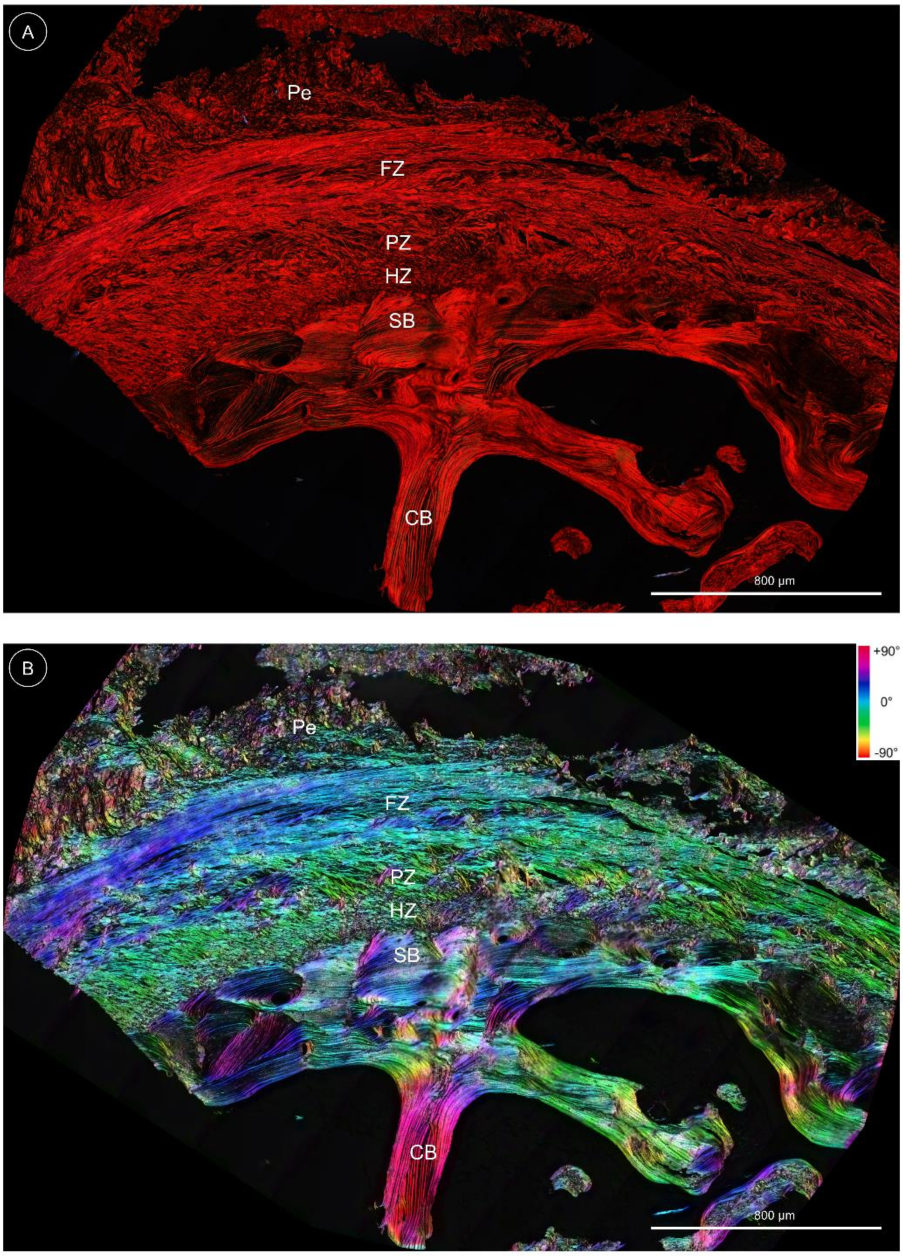

**Supplemental Figure 10. Polarized light microscopy images of the human mandibular condyle.** (A) Original polarized light microscopy image. (B) Color-coded image processed with OrientationJ in FIJI.

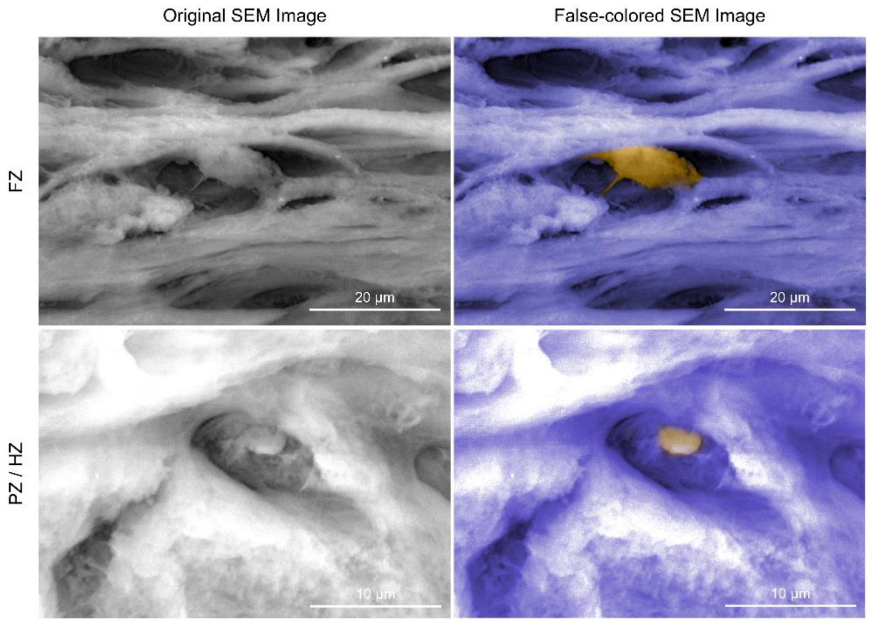


**Supplemental Figure 11. High-magnification scanning electron microscopy images of cells in guided engineered tissue.** Two distinct cell morphologies were observed: one in the fibrous zone (FZ) and another in the proliferative and hyaline zones (PZ/HZ).

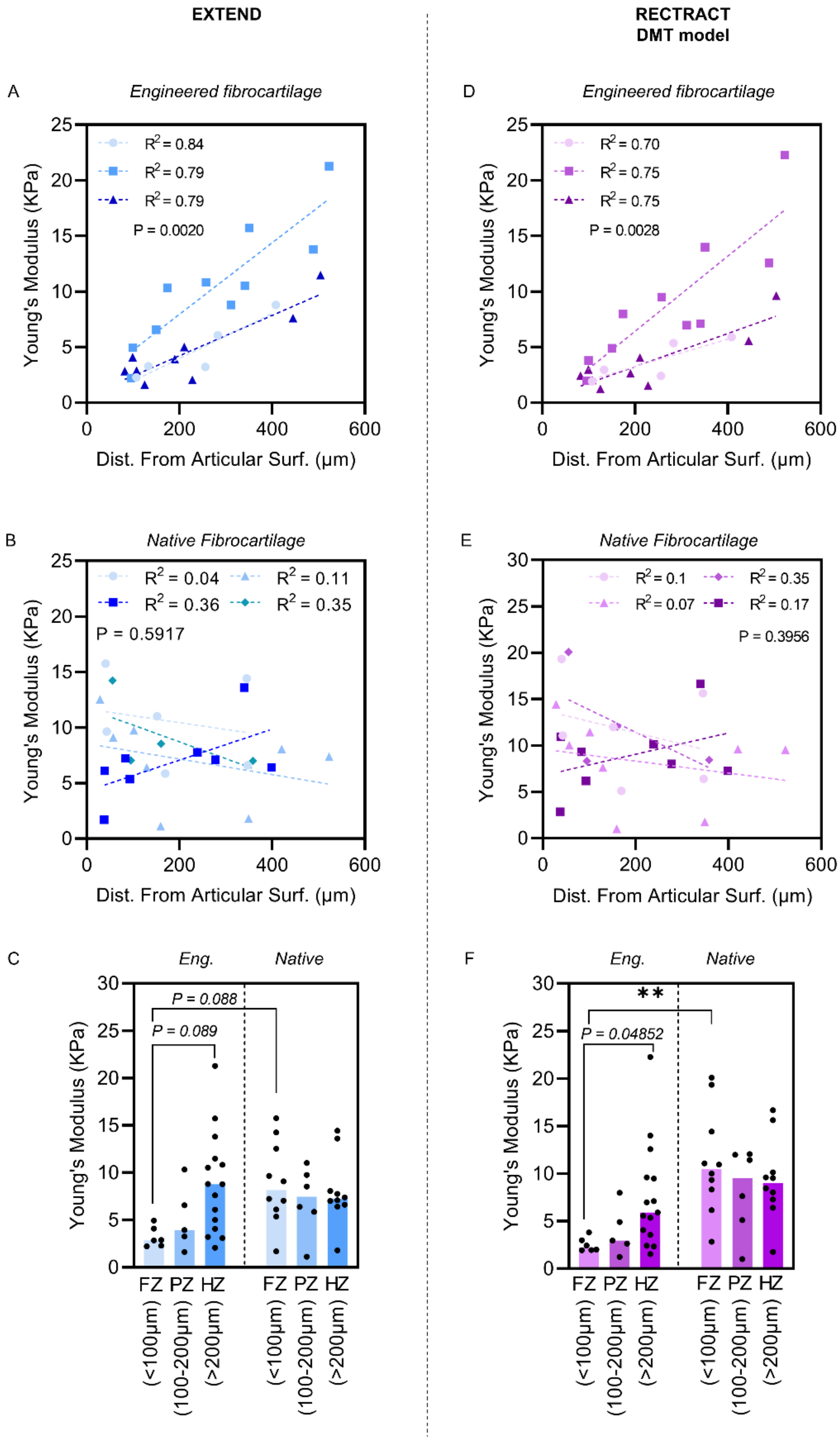


**Supplemental Figure 12. Young's modulus from extension and retraction curves**. The extension curve was processed using the standard Hertz model, while the DMT model accounted for solid adhesion in the retraction curve. Atomic force microscopy was performed on (A, D) guided engineered and (B, E) native cartilage cryosections in a depth-dependent manner to determine zonal micro-mechanical properties. (C, F) Young's modulus reported as zone-specific values.

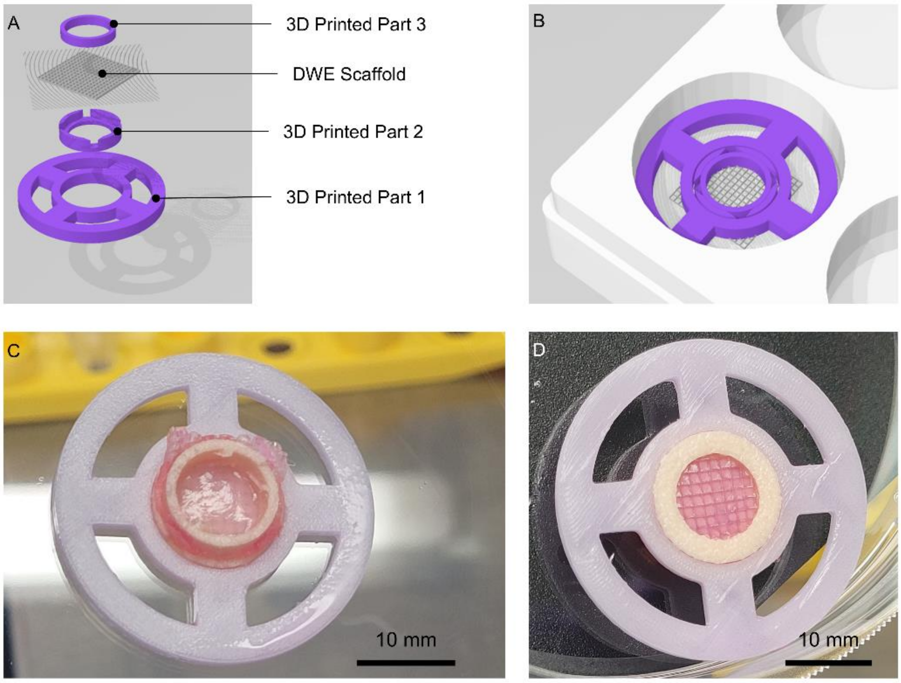


**Supplemental Figure 13**. **Insert system for positioning DWE scaffold and culturing guided engineered fibrocartilage**. (A) Four parts (purple) were 3D printed to restrict the seeding area to a cylinder with a diameter of 10 mm and a height of 1 mm (matching the DWE scaffold's height). (B) The system fitted into a 6-well plate. (C) Top view and (D) bottom view of the engineered guided fibrocartilage within the culture system.

**Supplemental Table 1.** Nuclei Distribution and morphology (mean ± std. dev.).

| | | *Nuclei / mm²* | *Circularity* | *Aspect ratio* |
|---|---|---|---|---|
| *Engineered tissue (unguided)* | FZ | 1,188 ± 287.4 | 0.7563 ± 0.1844 | 2.316 ± 1.054 |
| | PZ | 5,775 ± 1215 | 0.8382 ± 0.1575 | 1.712 ± 0.6405 |
| | HZ | 4,401 ± 607.7 | 0.8576 ± 0.1282 | 1.600 ± 0.4722 |
| *Engineered tissue (guided)* | FZ | 1,327 ± 468.7 | 0.8368 ± 0.1641 | 1.967 ± 0.9015 |
| | PZ | 4,985 ± 765.0 | 0.8865 ± 0.1367 | 1.616 ± 0.5736 |
| | HZ | 5,176 ± 797.5 | 0.8907 ± 0.1265 | 1.562 ± 0.5057 |
| *Human fibrocartilage (mandibular condyle)* | FZ | 735.0 ± 228.5 | 0.8184 ± 0.1680 | 1.999 ± 0.8602 |
| | PZ | 1,623 ± 478.6 | 0.8609 ± 0.1388 | 1.780 ± 0.6430 |
| | HZ | 618.6 ± 201.3 | 0.8890 ± 0.1179 | 1.683 ± 0.6485 |

**Supplemental Table 2.** Collagen fibril direction, dispersion and quality of alignment (mean ± std. dev.).

| | | *Direction (°)* | *Dispersion (°)* | *Quality of alignment* |
|---|---|---|---|---|
| *Engineered tissue (unguided)* | FZ | 21.1 ± 13.88 | 24.81 ± 3.816 | 0.9304 ± 0.1113 |
| | PZ | 28.82 ± 23.97 | 28.24 ± 9.005 | 0.8115 ± 0.1898 |
| | HZ | 38.47 ± 25.26 | 21.76 ± 6.637 | 0.7220 ± 0.2324 |
| *Engineered tissue (guided)* | FZ | 5.561 ± 5.782 | 23.01 ± 3.885 | 0.9828 ± 0.01571 |
| | PZ | 17.18 ± 16.30 | 31.29 ± 10.61 | 0.8083 ± 0.1684 |
| | HZ | 40.23 ± 27.92 | 28.99 ± 16.08 | 0.5686 ± 0.2319 |
| *Human fibrocartilage (mandibular condyle)* | FZ | 5.831 ± 4.704 | 12.43 ± 4.399 | 0.9233 ± 0.04444 |
| | PZ | 18.68 ± 9.876 | 20.78 ± 9.204 | 0.8373 ± 0.1444 |
| | HZ | 37.6 ± 13.25 | 22.19 ± 8.125 | 0.7573 ± 0.1365 |

**Supplemental Table 3.** Values of Young's modulus. adhesion force. and energy dissipation obtained from AFM experiments.

| | | Engineered tissue (guided) | | | Human fibrocartilage (mandibular condyle) | | |
|---|---|---|---|---|---|---|---|
| | | FZ | PZ | HZ | FZ | PZ | HZ |
| Young's modulus (KPa) *Extend Curve Hertz model* | Mean | 3.206 | 5.134 | 8.818 | 8.868 | 7.111 | 8.014 |
| | Std. Dev. | 1.082 | 3.403 | 5.329 | 4.317 | 3.529 | 3.617 |
| | Median | 2.856 | 3.930 | 8.787 | 8.161 | 7.463 | 7.236 |
| | Range | 2.714 | 8.717 | 19.22 | 14.06 | 9.904 | 12.63 |
| Young's modulus (KPa) *Retract Curve Hertz model* | Mean | 4.182 | 6.454 | 10.39 | 13.01 | 8.806 | 10.18 |
| | Std. Dev. | 1.638 | 4.411 | 5.388 | 6.813 | 4.425 | 4.953 |
| | Median | 3.964 | 5.471 | 10.10 | 10.46 | 9.640 | 9.399 |
| | Range | 4.348 | 10.70 | 16.50 | 22.77 | 12.31 | 17.92 |
| Young's modulus (KPa) *Retract Curve DMT model* | Mean | 2.522 | 3.939 | 7.515 | 11.26 | 8.204 | 9.348 |
| | Std. Dev. | 0.7449 | 2.605 | 5.488 | 5.416 | 4.495 | 4.302 |
| | Median | 2.224 | 2.971 | 5.901 | 10.49 | 9.534 | 8.995 |
| | Range | 1.869 | 6.735 | 20.74 | 17.24 | 11.02 | 14.89 |
| Adhesion Force (nN) | Mean | 4.030 | 3.125 | 1.709 | 0.9574 | 0.8053 | 0.9588 |
| | Std. Dev. | 2.258 | 2.732 | 1.797 | 0.5775 | 0.5003 | 0.5037 |
| | Median | 3.342 | 1.200 | 0.9587 | 0.7794 | 0.6343 | 0.9412 |
| | Range | 5.869 | 5.238 | 5.170 | 1.954 | 1.256 | 1.500 |
| Energy Dissipation (fJ) | Mean | 6.953 | 5.734 | 3.279 | 1.636 | 1.302 | 1.723 |
| | Std. Dev. | 3.760 | 4.850 | 3.212 | 0.9414 | 0.7403 | 0.9032 |
| | Median | 6.031 | 2.758 | 1.651 | 1.311 | 1.299 | 1.399 |
| | Range | 8.571 | 9.989 | 9.354 | 2.946 | 2.266 | 3.194 |
